# Prenatal Exposure to Bacterial Extracellular Vesicles Influences Fetal Gut Immunity and Immune Programming

**DOI:** 10.64898/2026.01.15.699706

**Authors:** Manuel S. Vidal, Ananth Kumar Kammala, Madhuri Tatiparthy, Ryan C. V. Lintao, Rahul Cherukuri, Ourlad Azeleus Tantengco, Shelly A Buffington, Enkhtuya Radnaa, Lauren S Richardson, Ramkumar Menon

## Abstract

**Background:** The fetal immune system undergoes pivotal development during gestation, preparing for postnatal antigenic challenges. Bacterial extracellular vesicles (bEVs), bioactive particles shed by bacteria, are emerging as modulators of host immunity. However, their role in shaping fetal intestinal immune development remains largely unexplored.

**Objectives:** This study aimed to investigate the effects of bEV exposure on lymphoid and myeloid populations in the fetal murine gut, focusing on their role in priming intestinal immunity, promoting differentiation, and modulating immune cell phenotypes in both normal and germ-free (GF) environments.

**Materials and Methods:** We used a murine model to evaluate the immune-modulating effects of bEVs during fetal development. bEVs were isolated from bacterial cultures and introduced into the amniotic sac of embryonic day 15.5 (E15.5) fetuses through intra-amniotic injection. Fetal and neonatal mice were either raised under conventional conditions (normal environment, NE) or in germ-free (GF) environments to assess microbiota-dependent effects. Immune profiling of fetal (E17) and postnatal (4 weeks) gut tissues was performed using high-dimensional mass cytometry (CyTOF) in both conventionally housed and germ-free (GF) mice. Clustering and differential expression analyses identified lymphoid and myeloid subpopulations, including progenitors, antigen-presenting cells, and intestinal stem cells (ISCs). secondary immune challenge (LPS or TSST-1) was conducted in postnatal bEV-primed mice to assess immune memory responses.

**Results:** bEV exposure significantly increased the prevalence of CD45- CD24+ CD44+ ISCs, promoting intestinal renewal and defense via differentiation into Paneth and tuft cells. These ISCs exhibited potential antigen-presenting capabilities through MHC expression. CD45+ lymphoid progenitors were upregulated, highlighting their role in early differentiation pathways. Myeloid progenitors, particularly monocyte-dendritic progenitor subsets, showed a bias toward antigen-presenting phenotypes.Germ-free models revealed heightened sensitivity to bEVs, with pronounced activation of progenitors and a reduction in exhaustion markers. Interestingly, macrophage and neutrophil populations displayed dose-dependent modulation, with low bEV concentrations promoting their expansion and higher doses leading to reduced incidence. Our findings suggest that bEVs act as immune priming agents in the fetal gut, promoting progenitor expansion and differentiation while preparing the intestine for postnatal challenges. Differences in responses between NE and GF models emphasize the importance of environmental influences, including microbiota, on bEV-mediated immune modulation.

**Conclusion:** bEVs play a pivotal role in shaping fetal intestinal immunity by priming lymphoid and myeloid progenitors and enhancing ISC function. These results open potential avenues for leveraging bEVs in immunomodulation and vaccine strategies. Future studies should explore the functional responses of bEV-primed cells and their translational relevance in humans.

## 1. Introduction

The regulatory factors and mechanisms that control immune system development and function are highly complex. Human immune system development begins *in utero*. Programming of the fetal immune system occurs at discrete stages of gestation and is known to be influenced by both maternal and fetal signaling. However, this is an understudied area primarily due to the difficulty of carefully accessing the human fetus during pregnancy. Current research on reproductive immunology has focused on the tolerogenic mechanisms at the feto-placental-maternal interface that allow pregnancy maintenance^1–12^. Protective immunity is the arm of reproductive immunology concerned with how progressive education of the fetal immune system programs its maturation *in utero*^13–16^. Protective mechanisms, like early training and education of the fetal immune system, are of interest as this system must produce existential responses to rapid microbial and other antigenic challenges *ex utero*^17–19^. Consequently, *in utero* education of the immune system and its imprinting in response to exposures during pregnancy (*e.g.*, maternal vaccination) is fundamental to priming of the emerging human immune system^20–25^. Among exposures, maternal inflammation during pregnancy has been shown to program long-term immune output from hematopoietic progenitors and subsequently impact offspring immune function in mouse models^26^; however, the precise mechanisms by which this programming occurs are unknown.

Extracellular vesicles (EVs; exosomes, 30-200nm size particles) are one of the most dynamic products that cells produce throughout their lifespan^27^. EVs were once considered carriers of cellular metabolic waste; however, emerging data demonstrate that their roles in various cellular biological functions are vast and mostly unknown^28^. During pregnancy, feto-maternal paracrine communication by EVs is one of the fundamental bases of maintaining homeostasis. EVs also signal parturition at term and preterm pregnancies^29,30^. Recently, we reported the discovery of bacterial extracellular vesicles (bEVs) released from commensal microbes of the host (various maternal body sites) and their presence in the placenta. bEVs are also reported in the amniotic fluid of pregnant women during normal pregnancy^31^. As reported, bEVs can elicit an immune response in host macrophages at supraphysiologic doses. While bEVs in the placenta are not proinflammatory, they do generate an immune response in the fetus.

It is postulated that exosomes can interact with various immune cells in the body. In innate immunity, nucleic acids can be packaged inside exosomes derived from T-cells, virus-infected cells, and cancer cells to be subsequently targeted by dendritic cells for presentation and interferon release^32–35^. Macrophages are also able to be either polarized or inactivated by tumor-associated exosomes *via* the delivery of non-coding RNAs and proteins^36–39^. Neutrophil extracellular trap release and interleukin production from neutrophils have also been observed to be abrogated *via* exposure to exosomes^40^. In adaptive immunity, plasma exosomes have been shown to activate T-cells, possibly through packaged MHCs inside exosomes of dendritic cells^41,42^, while also transforming the T-cells into a suppressive phenotype *via* interactions with multiple tumor-derived exosomes^43^.

During pregnancy, EVs coming from various body sources can be deposited in the placenta and traverse the placenta to allow exposure to the growing fetus and affect the physiology of fetal tissues^29,30,44^. Little is known about the propagation and effect of bEVs on fetal tissues. In a cellular environment where the majority of immune cells are undergoing training and development, bEV exposure might alter the differentiation of specific precursor cell populations toward certain phenotypes. Here, we reasoned that training may not necessarily be induced *via* inflammation, as overwhelming inflammation is likely to be deleterious for the pregnancy; instead, a fine-tuned balance of stimulation would be a more favorable environment for appropriate fetal immune development. We specifically tested the hypothesis that bEV from maternal body sites systemically deposited in the placenta can reach the fetus, causing *in utero* immune education and rendering the neonatal immune system programmed to recognize these microbes as self.

In this study, we used intraamniotic injections of bEVs in pregnant mice and high-dimensional mass cytometry to investigate how bEVs affect the development of lymphoid and myeloid immune cell populations in fetal and postnatal gut tissues. The results reveal that bEV exposure induces a dose-dependent upregulation of diverse immune —including progenitors, activated lymphocytes, and dendritic-like cells,with distinct patterns observed between mice raised in a conventional (normal-) environment and those raised in a reduced-germ load environment as well as between embryonic and young mice. Overall, our findings suggest that bEVs can prime and educate the developing immune system, offering promising insights into potential novel early-life immunization strategies based on extracellular vesicles.

## 2. Materials and Methods

### 2.1 Institutional ethics approval

All Animal procedures were followed in accordance with the Institutional Animal Care and Use Committee (IACUC) at the University of Texas Medical Branch, Galveston, under approved protocol number 041107F.

### 2.2 Husbandry and mating

Normal-environment (NE) CD-1 mice were maintained at 4–5 mice/cage in standard cages in the UTMB Animal Research Center housing facility under a strict 12-h light/dark cycle. Normal chow (Lab Diets, #5001) and osmotically-sterilized drinking water were provided *ad libitum*.

Germ-free (GF) C57BL/6 mice were kindly provided by Dr. Richard Pyles’ laboratory and maintained in a separate building under a reduced-germ load environment. Autoclaved chow and water were provided to these mice *ad libitum*. Nulligravid mice were mated with male mice (n=3 for breeding) and, upon confirmation of a vaginal plug (designated as embryonic day 1, E1), the bred dams were placed in individual cages until E15.

### 2.3 Protocol for the isolation of bEVs from the *E. coli* and human placenta

bEVs were isolated following the established protocol from Menon *et al*.^31^ *Escherichia* (*E*.) *coli* ATCC 12014 strain O55:K59(B5):H, obtained from Remel Laboratory of Thermo Fisher (Thermo Fisher Scientific, Remel Products, Lenexa, KS, United States, Lot # 496291), was cultured in nutrient broth (BD Biosciences) with stocks stored in 20% glycerol at −80°C. Logarithmic phase cultures were processed using the ExoBacteria™ OMV Isolation Kit (Cat# EXOBAC100A-1, System Biosciences, Palo Alto, CA, United States) to isolate bEVs. Placental specimens for human placenta bEV isolation were deidentified and considered as discarded human specimens that do not require institutional review board (IRB) approval. Placental specimens were collected from John Sealy Hospital at the University of Texas Medical Branch at Galveston, Texas, USA, in accordance with the relevant guidelines and regulations of approved protocols for various studies (UTMB 11-251; University of Texas Medical Branch at Galveston). Specimens were first processed according to the established protocol from Vidal *et al*.^45^ Briefly, tissues were minced into uniform 2 mm³ pieces, digested in endotoxin-free PBS containing collagenase D (2 mg/mL) and DNAse I (40 U/mL) at 37°C with constant rotation, and then filtered and centrifuged to obtain a crude extract, which was concentrated using 10-kDa Amicon tubes. This concentrated extract was layered into an iodixanol density gradient (comprising 50%, 40%, 20%, and 10% iodixanol solutions topped with PBS) and subjected to ultracentrifugation at 100,000×*g* for 18 hours at 4°C, after which the 8^th^ and 9^th^ fractions—containing the placental bEVs—were carefully collected on ice. All bEVs were stored at −80°C prior to use.

### 2.4 Induction of bEV exposure in pregnant mice

At E15, the pregnant mice were brought to our Animal Surgery Room. Each mouse was anesthetized with 1-2% isoflurane in oxygen delivered through a nosecone using a controlled-delivery anesthetic machine. Once unconsciousness was determined, individual mice were placed supine in a sterile surgical field. Limb restraints were put into place. A midline abdominal incision was performed, and the uterine horns were exposed. A single intraamniotic injection of each treatment [10 µL of one of the following: *E. coli*-derived bEVs (9 × 10^6^ particles) (EC-bEV), “low-dose” human placental bEVs (9 × 10^6^ particles), “high-dose” human placental bEVs (9 × 10^8^ particles), or endotoxin-free PBS] was administered per amniotic sac. After injections had been completed, the uterine horns were repositioned to their original location, the abdominal walls were positioned, and the defect was closed using sterile surgical staples. A single subcutaneous injection of buprenorphine (0.05 mg/kg/dose) was given to each mouse. The pregnant mice were transferred back to clean individual cages. Heat lamps were employed to warm the mice during recovery. Monitoring was performed every 2 hours, and any signs of weakness were accounted for. Mice were transferred to the Animal Satellite Room to recover. The experimental design is shown in Figure 1.

**Figure 1.**
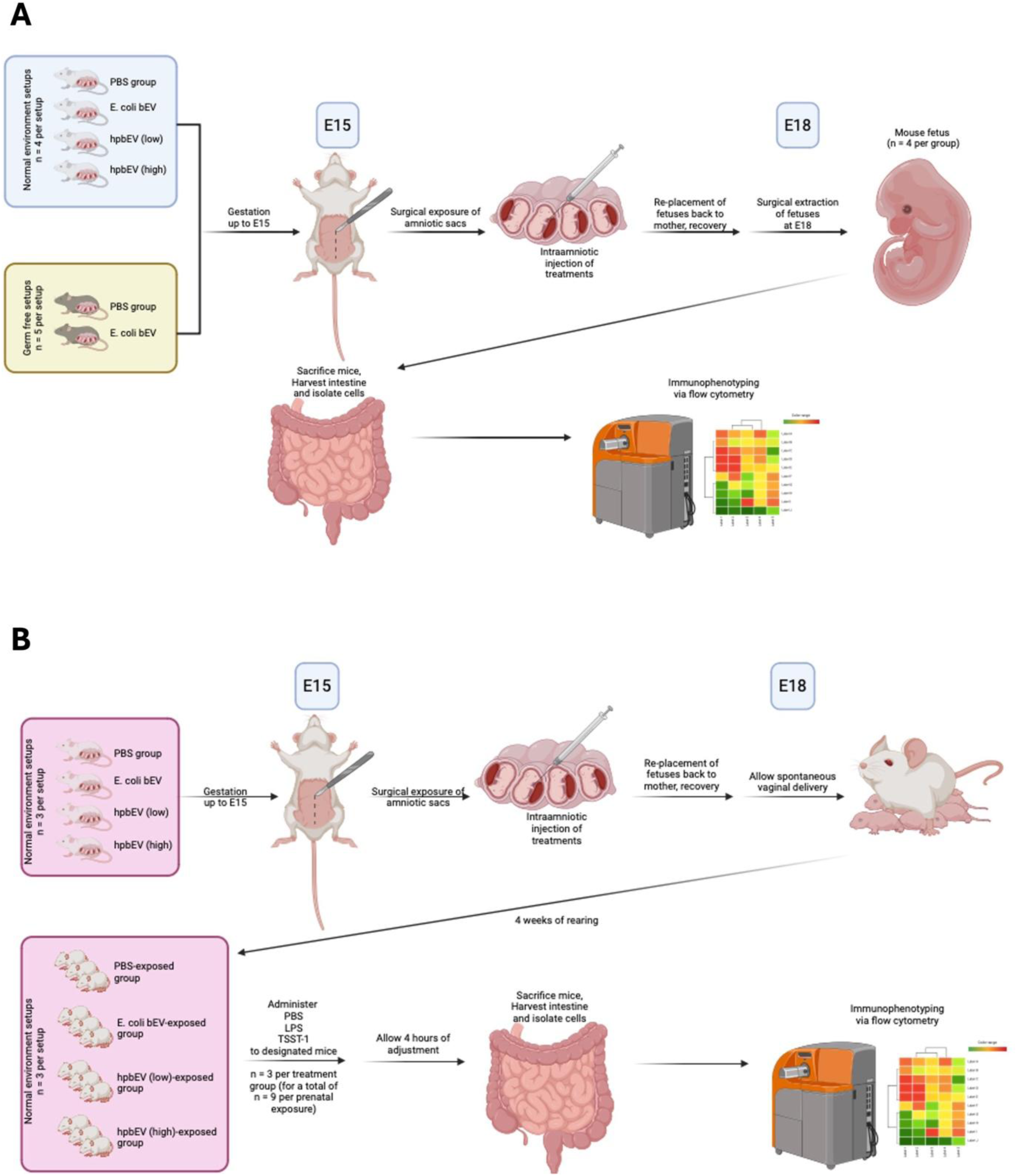
Experimental design for the study. (A) Flow for the first part of the study, exploring cell populations during fetal life. (B) Flow for the second part of the study, exploring early responses to bacterial extracellular vesicle (bEV)-primed mice. PBS, phosphate-buffered saline. hpbEV, human placenta bEV. LPS, lipopolysaccharide. TSST-1, toxic syndrome shock toxin-1.

#### 2.4.1 Tissue harvesting

On E17, the treated pregnant mice were transferred again to our Animal Surgery Room. The mice were euthanized *via* carbon dioxide asphyxiation. Once death was confirmed, mice were placed supine in a sterile surgical field. A midline abdominal incision was performed, and the uterine horns were exposed. Incisions were made per amniotic sac, and fetuses were harvested. Each pup’s intestines were individually obtained and stored in RPMI media with 10% FBS and 1% penicillin-streptomycin for cell isolation.

### 2.5 bEV priming and postnatal immune challenge

NE CD-1 mice were assigned to three groups. Each of the groups were given different “priming” exposures – the first batch was given an intraamniotic injection of endotoxin-free PBS, the second batch was given EC-bEV (9 × 10^5^ particles), and the third batch was given LD-bEV (9 × 10^5^ particles) (Figure 1). The doses for this priming challenge were predetermined by exposing GF mice to various doses of EC-bEV, and the lowest dose, which did not cause mortality, was chosen. The administration method was similar to the aforementioned surgical technique. The animals were allowed to deliver spontaneously, and the pups were kept with the mother until four weeks of age. At this point, three young mice from each group were then separated accordingly and given separate treatments *via* oral gavage – one group was given endotoxin-free PBS, one group was given lipopolysaccharide (50 mg/kg), and one group was given toxin shock syndrome toxin-1 (TSST-1) (0.01 ug/kg). The mice were then allowed to recover, and, four hours later, they were individually euthanized. Embryonic pup intestines were harvested according to the protocol as mentioned earlier. All isolated tissues were kept at −80°C until cell isolation was performed.

#### 2.5.1 Isolation of cells from tissues

Upon collection of all necessary tissues, 500 µL of accutase was added, and the tissues were homogenized. The homogenates were then incubated at 37°C with shaking for 1 hr. Another 500 µL of RPMI was added to the homogenates, which were strained using a 70 µm cell strainer. The strained suspension was washed with repeated additions of RPMI and centrifugation at 1500 rpm for 5 min at 20°C. After the washing, the supernatant was removed, and the pellet was resuspended in 1 mL of RBC lysis buffer. After incubation for 10 mins, the suspension was neutralized with 9 mL of DMEM-F12/10% FBS and spun down at 1500 rpm for 5 min at 20°C. The supernatant was removed again, and 500 µL of cell freezing media was added to allow for storage at −80°C until staining.

### 2.6 High-dimensional single-cell profiling of feto-maternal tissues by mass cytometry (CyTOF)

#### 2.6.1 Antibodies

The CyTOF panel was designed based on available literature for known lymphoid and myeloid populations. A summary of antibodies used for each panel can be found in Table 1. The antibodies were sourced from the MD Anderson Cancer Research Center Flow core facilities. (MDACC, Texas, Houston), or custom conjugated using the Maxpar antibody conjugation kit (Fluidigm, Markham, ON, Canada) following the manufacturer’s protocol. After being labeled with their corresponding metal conjugate, the percentage yield was determined by measuring their absorbance at 280 nm using a Nanodrop 2000 spectrophotometer (Thermo Scientific, Wilmington, DE). Antibodies were diluted using Candor phosphate-buffered saline (PBS) antibody stabilization solution (Candor Bioscience GmbH, Wangen, Germany) to 0.3 mg/mL and then stored at 4°C. A summary of the marker antibodies is shown in Table 1.

**Table 1.** List of conjugated antibodies used for both lymphoid and myeloid panels, respectively.

| LYMPHOID |  | MYELOID |  |
| --- | --- | --- | --- |
| Target | Label | Target | Label |
| CD69 | 156Gd | Target | label |
| CD27 | 148Nd | CD11c | 209Bi |
| CD183, CXCR3 | 143Nd | I-A/I-E, MHC-II | 115In |
| CD62L | 164Dy | Ly-6G/C, Gr-1 | 141Pr |
| CD44 | 153Eu | CD11b | 143Nd |
| CD4(Ms) | 115In | CD14 | 156Gd |
| CD184, CXCR4 | 159Tb | FceR1a | 144Nd |
| CD197, CCR7 | 155Gd | CD117, c-kit | 166Er |
| CD24 | 169Tm | XCR1, GPR5, CCXCR1 | 168Er |
| CD279, PD1 | 165Ho | CD206, MMR | 169Tm |
| CD25 | 167Er | F4/80 | 174Yb |
| NK1.1, CD161b/c, Ly-55 | 170Er | CD86 | 172Yb |
| Granzyme B | 173Yb | CD8a | 146Nd |
| CD38 | 175Lu | Siglec-F | 172Yb |
| CD8a | 168Er | Ly-6C | 150Nd |
| CD45(Ms) | 89Y | Arg-1, ARG1 | 164Dy |
| TCRgd | 160Gd | CD103 | 147Sm |
| CD19 | 149Sm |  |  |
| Foxp3 | 158Gd |  |  |
| GATA3 | 145Nd |  |  |
| RORg(t) | 163Dy |  |  |
| T-bet | 161Dy |  |  |

#### 2.6.2 Antibody staining

Single-cell suspension samples were resuspended in Maxpar staining buffer for 10 min at room temperature on a shaker to block F_c_ receptors. Cells were mixed with a cocktail of metal-conjugated surface marker antibodies (Table 1), yielding 500-μL final reaction volumes, and stained at room temperature for 30 min on a shaker. Following staining, cells were washed twice with PBS with 0.5% BSA and 0.02% NaN_3_. Next, cells were permeabilized with 4°C methanol for 10 min at 4 °C. Cells were then washed twice in PBS with 0.5% BSA and 0.02% NaN_3_ to remove the remaining methanol. They were stained with intracellular antibodies in 500 μL buffer for 30 min at room temperature on a shaker. Samples were then washed twice in PBS with 0.5% BSA and 0.02% NaN_3_. Cells were incubated overnight at 4°C with 1 mL of 1:4,000 191/193Ir DNA intercalator (Standard BioTools, Inc., Markham, ON) diluted in Maxpar fix/perm overnight. The following day, cells were washed once with PBS with 0.5% BSA and 0.02% NaN_3_ and then two times with double-deionized water.

#### 2.6.3 Mass cytometry

Prior to analysis, the stained and intercalated cell pellet was resuspended in ddH_2_O containing polystyrene normalization beads containing lanthanum-139, praseodymium-141, terbium-159, thulium-169, and lutetium-175 as described previously^46^. Stained cells were analyzed on a CyTOF 2 (Standard BioTools Inc., Markham, ON) outfitted with a Super Sampler sample introduction system (Victorian Airship & Scientific Apparatus, Alamo, CA) at an event rate of 200-to-300 cells per second. All mass cytometry files were normalized using the mass cytometry data normalization algorithm freely available for download from https://github.com/nolanlab/bead-normalization.

#### 2.6.4 Data Analysis

CyTOF data sets were first manually gated using Standard BioTools/Fluidigm clean-up procedure, including Gaussian discrimination (Markham, ON) in FlowJo V10 (FlowJo LLC). Then, each sample was given a unique Sample ID, then all samples were concatenated into a single.fcs file. This concatenated file was further analyzed by t-distributed stochastic neighbor embedding (t-SNE) in FlowJo V10, using equal numbers of cells from individual treatment groups. The analyses were done for both lymphoid and myeloid panels. tSNE was performed using the following settings: Iterations, 3000; Perplexity, 50; Eta (learning rate), 28000. Heatmaps of marker expression were generated using the Color Map Axis function. Visualizing the resulting t-SNE plot as a heatmap of marker expression of immune cells. To explore the phenotypic diversity of immune cell populations in the different groups of mice FM tissues, we applied a K-nearest-neighbor density-based clustering algorithm called Phenograph. This algorithm allows the unsupervised clustering analysis of data from single cells. The output was organized using the Cluster Explorer tool to visualize the phenotypic continuum of cell populations. This tool creates an interactive cluster Profile graph and heatmap and displays the cluster populations on a tSNE plot. For each cluster, marker positivity was set at ≥10x the relative expression level of the marker with the lowest expression level. Two-way ANOVA was employed to determine statistical differences between the groups; a p-level of <0.05 was considered significant.

## 3. RESULTS

### 3.1 bEV introduction drives distinct immune cell differentiation programs

First, to determine whether bEV introduction will stimulate mucosal fetal immunity, we introduced bEV particles mid-gestation intraamniotically and analyzed populations within the immediate term period. Using mass cytometry *via* time-of-flight (CyTOF) in isolated cells from murine gut tissues, we observed variably expressed clusters in the NE CD-1 mice in both lymphoid (35 clusters) and myeloid (21 clusters) populations. The delineated clusters with their corresponding markers are outlined in Tables 2 and 3. tSNE maps of lymphoid and myeloid populations in this setup are shown in Figure 2. Interestingly, we see an increase in the intensity of the populations identified, implying a general upregulation of cell populations upon bEV exposure.

**Figure 2.**
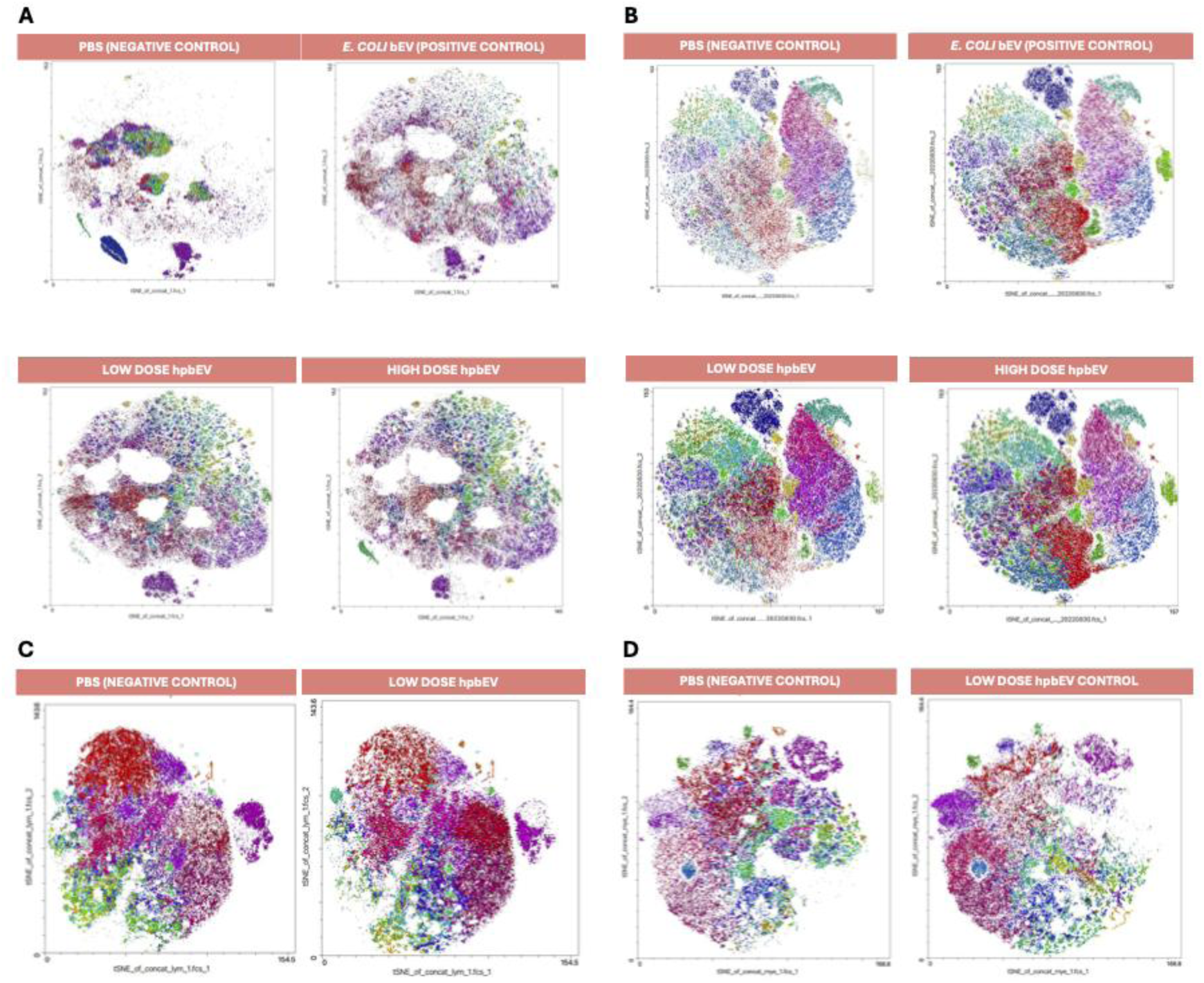
tSNE maps for lymphoid (first panel) and myeloid (second panel) populations in the normal environment setups (A, B) and germ-free (C, D) mice setups. Each tSNE panel features populations that vary in number of absolute events across different treatment setups, as visually apparent in the myeloid panel. Absent or present populations also vary across different treatment setups, as visually apparent in the lymphoid panel. PBS, phosphate buffered saline. bEV, bacterial extracellular vesicles. hpbEV, human placental bacterial extracellular vesicles.

**Table 2.**
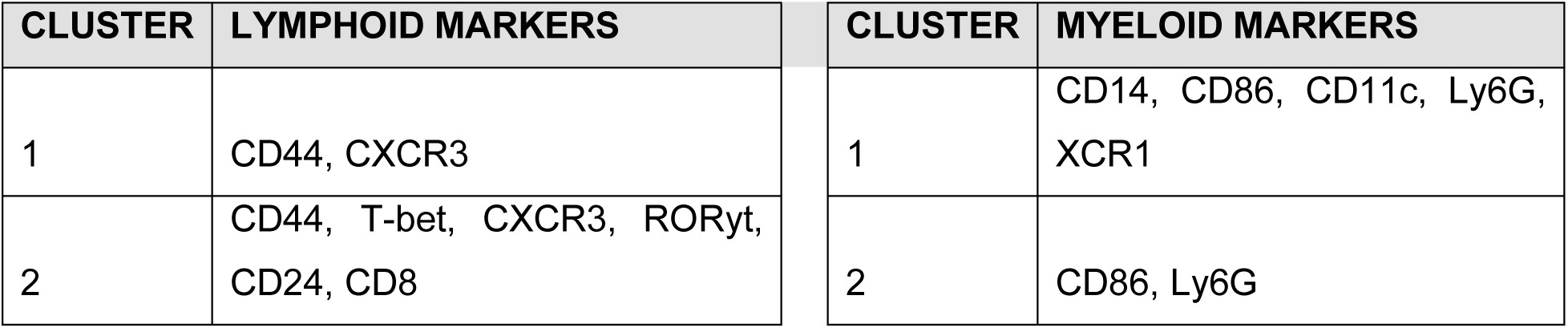
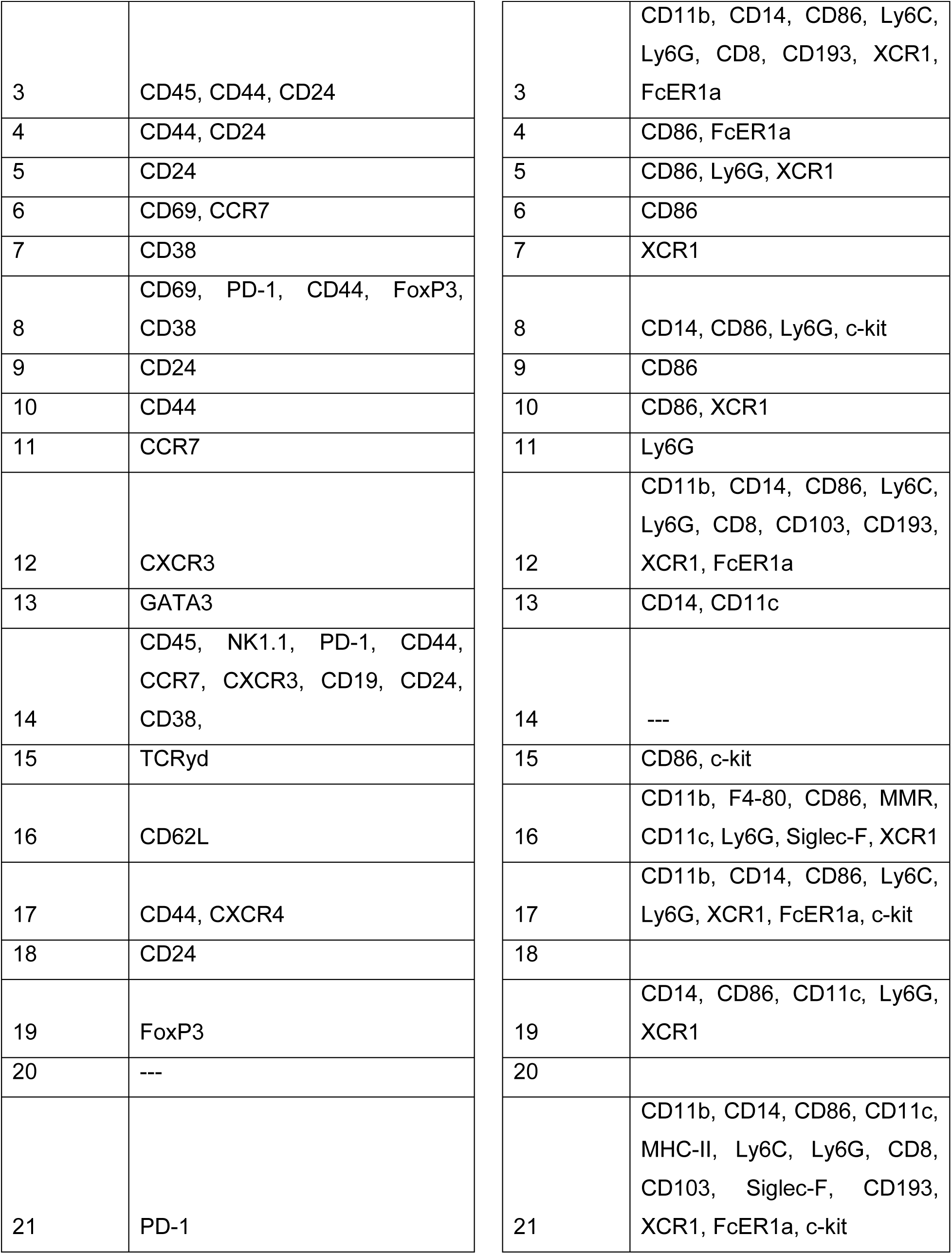
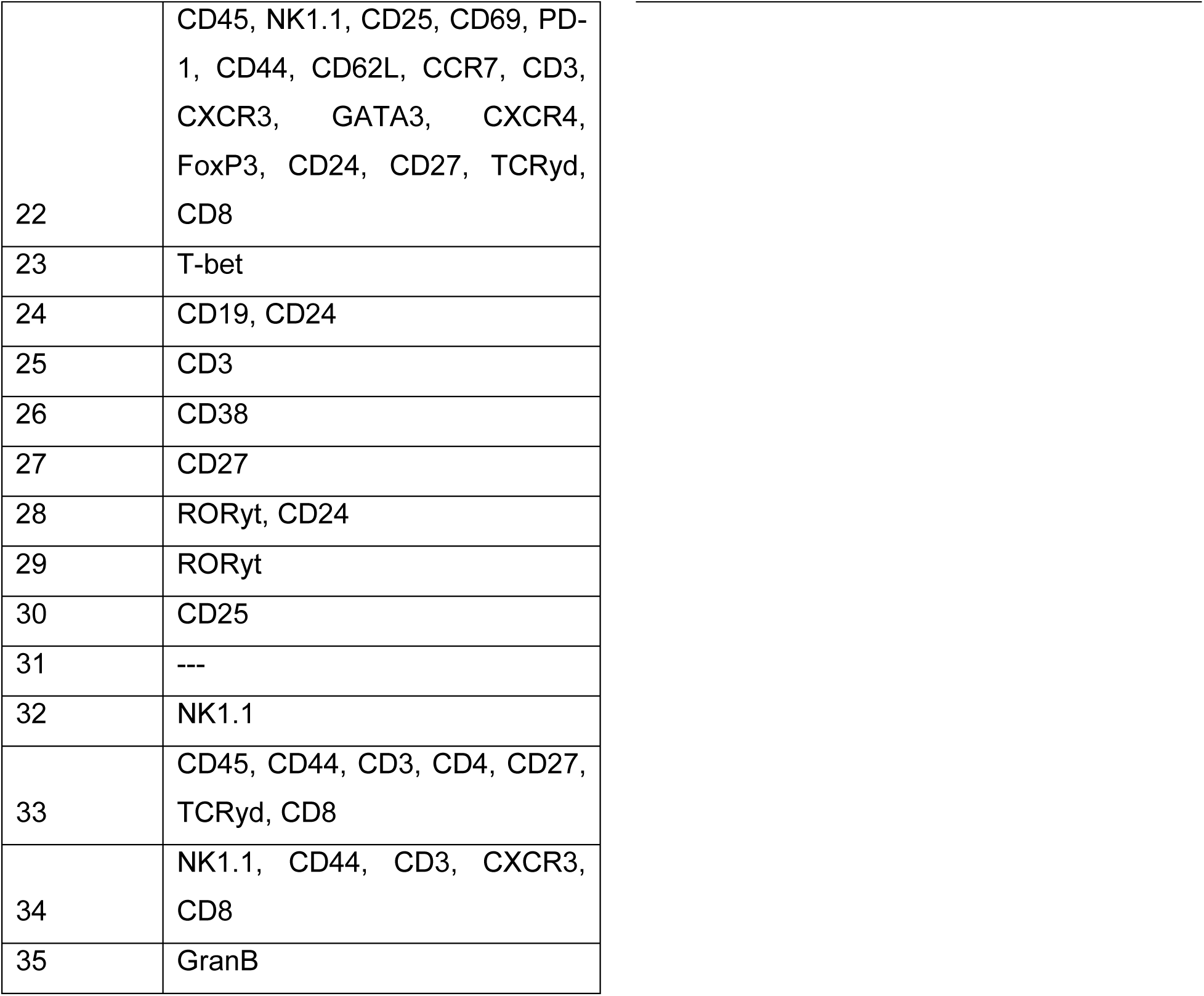
Cell clusters observed in the CyTOF analysis using a lymphoid and myeloid panel in normal-environment mice.

**Table 3.** Cell clusters observed in the CyTOF analysis using a lymphoid and myeloid panel in germ-free mice.

| CLUSTER | LYMPHOID MARKERS | CLUSTER | MYELOID MARKERS |
| --- | --- | --- | --- |
| 1 | CCR7, CD24 | 1 | CD14, Ly6G |
| 2 | CD69, CD44, CCR7, CXCR3 | 2 | Ly6G |
| 3 | CD45, CD44, CD24 | 3 | CD14, Ly6G, XCR1, FcER1a |
| 4 | CD44 | 4 | XCR1, FcER1a |
| 5 | CD69, CCR7, CXCR3 | 5 | FcER1a |
| 6 | CD69, PD-1, CD44, CD3, FOXP3, CD38 | 6 | CD14 |
| 7 | CD69, CD44, CCR7, CD24 | 7 | CD14, CD11c, Ly6C, Ly6G, CD8, CD103, XCR1, FcER1a, |
| 8 | CCR7 | 8 | CD14 |
| 9 | CD8, CD44 | 9 | c-kit |
| 10 | CD44 | 10 | XCR1 |
| 11 | CXCR3 | 11 | CD14, XCR1 |
| 12 | CD24 | 12 | Arg-1 |
| 13 | TCRγd | 13 | MMR, XCR1 |
| 14 | PD-1, GATA3 | 14 | CD11b |
| 15 | T-bet | 15 | CD193, XCR1 |
| 16 | CD62L, T-bet, RORγt | 16 | Ly6G, XCR1 |
| 17 | CD19, CD24 | 17 | CD86 |
| 18 | --- | 18 | XCR1 |
| 19 | CD45, CD69, CD44, CCR7, CXCR3, CD38 | 19 | Ly6G |
| 20 | CD44 | 20 | CD14, CD11c |
| 21 | FOXP3 | 21 | CD103 |
| 22 | CD62L | 22 | CD11b, F4-80, Ly6G, Siglec-F, FcεR1a |
| 23 | CXCR4 | 23 | Ly6C |
| 24 | --- | 24 | CD8 |
| 25 | CD86 | 25 | Ly6G |
| 26 | CD3 | 26 | XCR1 |
| 27 | --- | 27 |  |
| 28 | CD45, CD8, CD25, CD69, CCR7, CD4 | 28 | FcεR1a |
| 29 | CD45, NK1.1, CD8, CD25, CD69, PD-1, CD44, CD62L, CCR7, CD3, CD4, CXCR3, GATA-3, CXCR4, RORγt, FOXP3, CD19, CD24, CD27 | 29 | |

In Figures 3 and 4, various diverse populations with different markers display general upregulation upon bEV exposure. In the lymphoid subset, the majority of the cells are CD45-but express either of the memory markers CD44^47^, CCR7^48–50^, and CD62L^51–53^. The remaining CD45-clusters may represent populations of intestinal stem cells that can differentiate into epithelial cells, providing an initial barrier to pathogens and continually providing a pool of ready ISCs^54,55^. Additionally, we identified five CD45+ populations in the fetal gut of NG mice. One of the clusters (Cluster 29: CD45+ CD3+ CD8+ NK1.1+ CD25+ CD69+ PD-1+ CD44+ CD62L+ CCR7+ CXCR3+GATA3+ CXCR4+ FoxP3+ CD24+ CD27+ TCRyd+) may indicate a unique progenitor in fetal life. Another population, CD45+ CD24+ CD44+ cluster (Cluster 17), may point towards a unique set of lymphocytes, while the CD45+ CD4+ CD8+ population may point towards a set of double-positive T cells. Interestingly, two of the lymphoid clusters in NG mice are exhausted phenotypes (PD-1+) of NK cells (NK1.1+ CCR7+ CXCR3+ CD38+) (Clusters 32, 34) ^56–58^and regulatory T cell markers (FoxP3+ CD69+) (Cluster 8)^59,60^. Therefore, it may be presumed that in the normal environment, the majority of the lymphoid cells are poised for memory phenotypes, although early activation of certain populations is observable – an increase in progenitors, and a reduction in exhausted phenotype.

**Figure 3.**
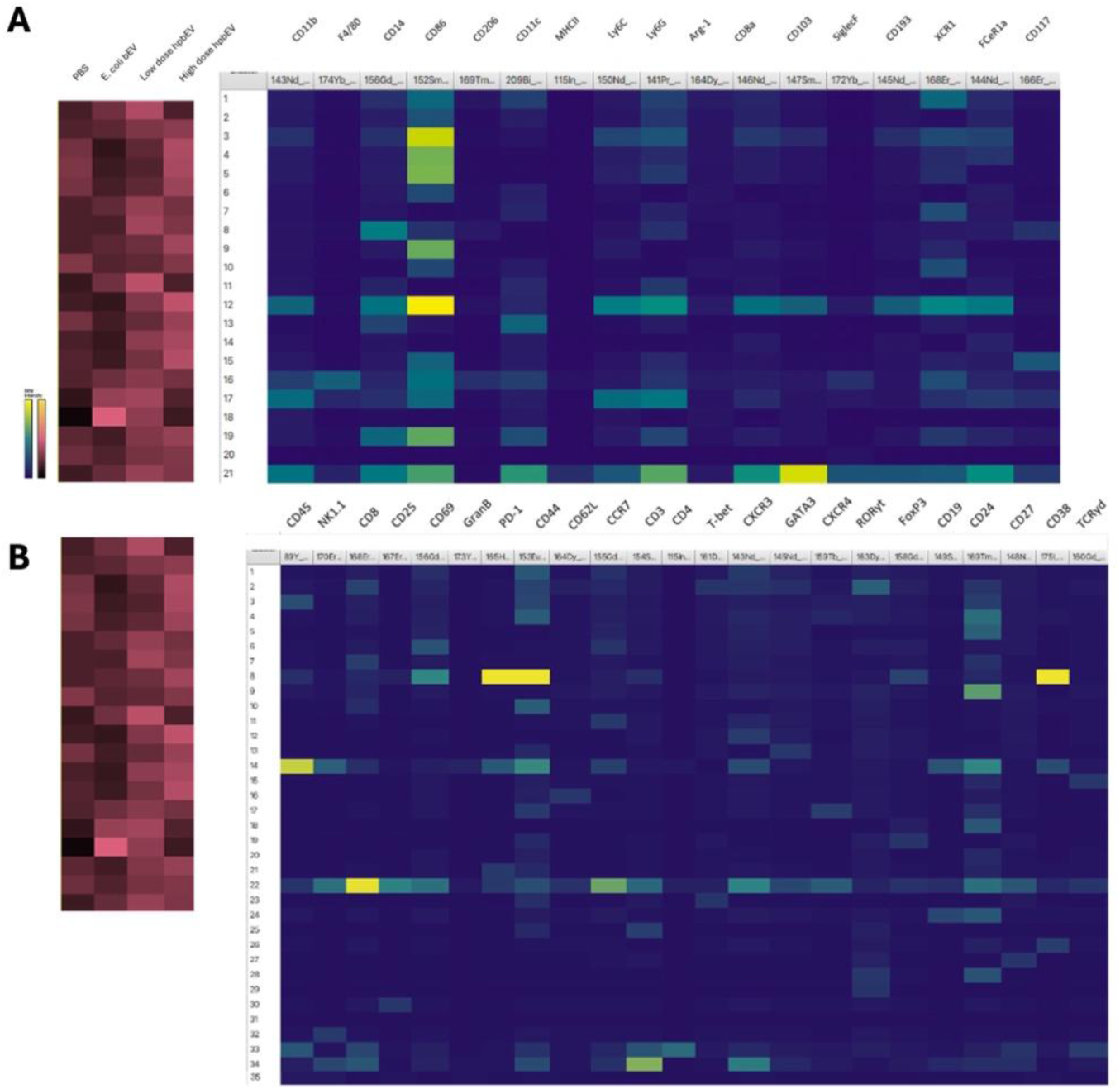
Heatmaps for the myeloid (A) and lymphoid (B) populations across different treatments in the normal environment mice setups. Each population can be broken down into different clusters, as shown in the heatmap (blue palette). Each cluster represents a specific cell type, whose marker positivities vary across each marker. Overall, 20 clusters were identified for the myeloid population and 35 clusters were identified for the lymphoid population. The relative abundances of the different clusters, or cell types, vary across different treatments, as shown in the heatmap (red palette). PBS, phosphate buffered saline. bEV, bacterial extracellular vesicles. hpbEV, human placental bacterial extracellular vesicles.

**Figure 4.**
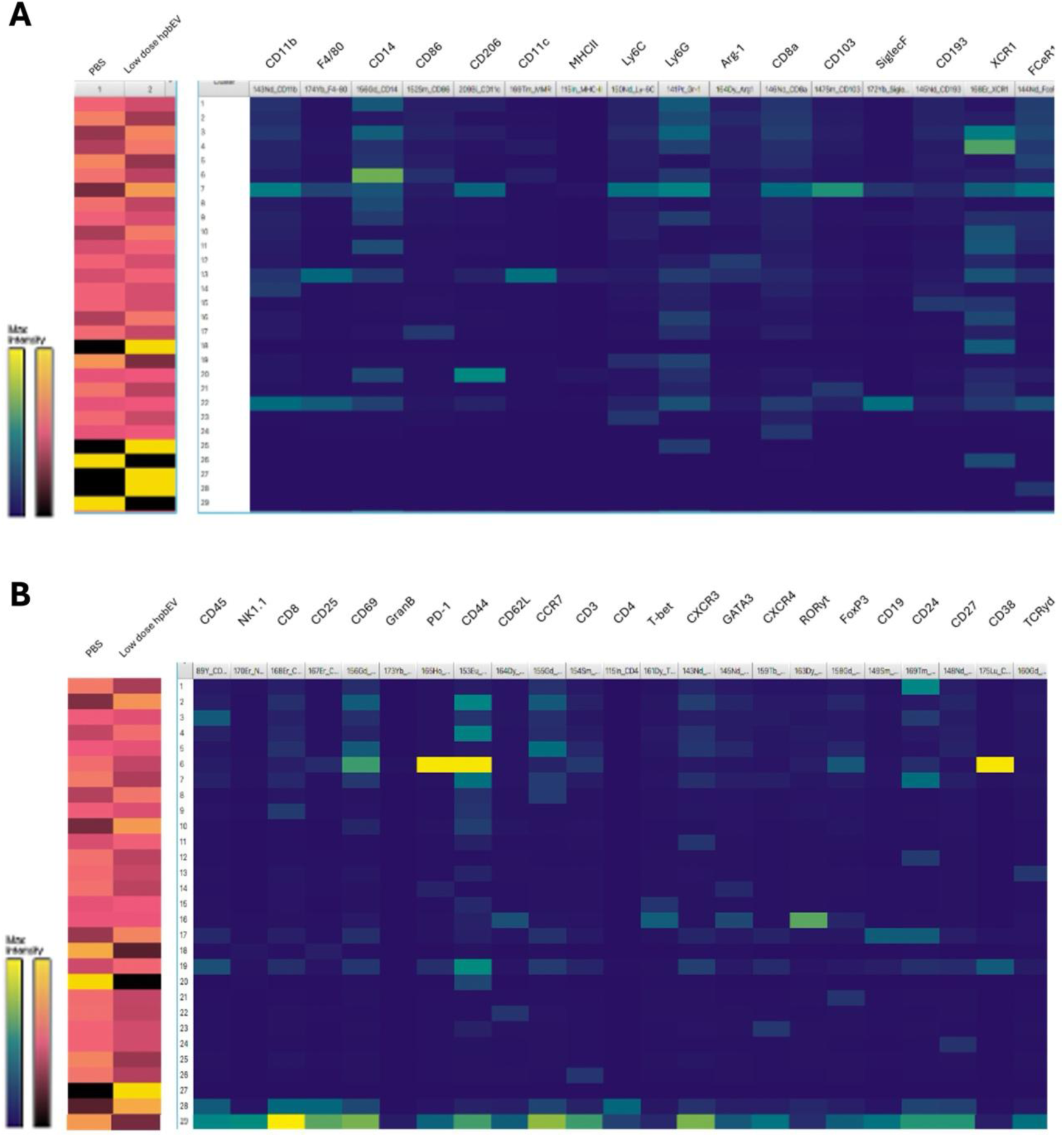
Heatmaps for the myeloid (A) and lymphoid (B) populations across different treatments in the germ-free mice setups. Each population can be broken down into different clusters, as shown in the heatmap (blue palette). Each cluster represents a specific cell type, whose marker positivities vary across each marker. Overall, 29 clusters were identified for the myeloid population and 29 clusters were identified for the lymphoid population. The relative abundances of the different clusters, or cell types, vary across different treatments, as shown in the heatmap (red palette). PBS, phosphate buffered saline. hpbEV, human placental bacterial extracellular vesicles.

In the myeloid subset, we observed clusters of CD11b+ cells, which are markers for monocyte/dendritic-like cells. We presume that four clusters (Cluster 3, 12, 17, 21) are putative monocyte-dendritic progenitors, since there were multiple monocyte/macrophage and dendritic cell markers that are present in PBS and high-dose bEV setups only. Another CD11b+ positive cluster (Cluster 16) was observed to increase only in low-dose bEV setups, which may represent intestinal macrophages (CD11b+ F4-80+ CD86+ CD11c+ XCR1+). Six clusters of CD11b-Ly-6g+ clusters are present (Clusters 1, 2, 5, 8, 11, 19), which in general increase with increasing bEV concentration. The third group of clusters is CD11b- Ly6G- CD86+ cells, which may be transient cells brought about by the gradual differentiation of progenitors into cells of monocyte and dendritic phenotype. Neutrophil-like Ly-6g+ clusters (Cluster 2, 5, 8, 11) increase proportionally to the bEV dose. Early response to bEVs, then, seems to be reliant on an increase in neutrophil-like cells, with potential differentiation of monocyte-dendritic progenitors into macrophage/dendritic-like cells.

### 3.2 Germ-free mice reveal activation of granulocytic-like cells and lymphoid progenitors in response to bEV stimulation

Second, to shed light on which immune cells will be activated in the mucosa of a bacterially-naïve system, we characterized the baseline state of gut immune cell populations in the embryonic gut of GF *versus* conventionally colonized mice with or without bEV treatment. Notably, due to a limited number of samples, we compared only a low-dose exposure of hpbEVs against a negative control in our GF setups. Overall, we observe differentially expressed lymphoid (29 clusters) and myeloid (29 clusters) populations in the pups of controls versus bEV-treated mice (side panels, Figure 4).

In the myeloid group (Figure 4A), there is an XCR1+ population (Cluster 18), Ly6G+ population (Cluster 25) and a FCeR1a+ population (Cluster 28) that are only present in the low-dose hpbEV, while there is an XCR1+ population (Cluster 26) that is only present in the PBS group. It appears that without the influence of native environment, neutrophil-like and eosinophil-like cells are the primary innate immune responses to bEV exposure, with varying populations of dendritic-like cells being activated concomitantly.

Interestingly, in the lymphoid group (Figure 4B), the majority of cell populations are at a higher incidence in the PBS group compared to the low-dose bEV group. Both groups express two multi-marked populations, although a more progenitor-like state is more prevalent in the PBS group (Cluster 29: CD45+ NK1.1+ CD8+ CD25+ CD69+ PD-1+ CD44+ CD62L+ CCR7+ CD3+ CD4+ CXCR3+ GATA-3+ CXCR4+ RORyt+ FOXP3+ CD19+ CD24+ CD27+) compared to the low-dose bEV group (Cluster 28: CD45+ CD8+ CD25+ CD69+ CCR7+ CD4+). A CD44+ population (Cluster 20) is also present in the PBS group but not in the low-dose bEV group. Four populations seem to be more activated upon bEV exposure – memory T-cell-like populations (Cluster 2: CD69+ CD44+ CCR7+ CXCR3+ and Cluster 19: CD45+ CD69+ CD44+ CCR7+CXCR3+ CD38+) and B-cell-like populations (Cluster 17: CD24+ CD44+ cells). Taken together, it appears that without the influence of the native environment, the lymphoid response upon bEV exposure shifts into an early memory response with concomitant humoral regulation.

### 3.3 Embryonic progenitors, mucosal macrophage-like populations, and dendritic-cell-like populations comprise the majority of the myeloid cell populations activated upon second encounter

Given that there is observable stimulation of fetal mucosal immunity upon bEV exposure, we theorized that (1) bEVs may contain bacterial antigens that can activate the immune system, (2) the initial encounter with bEVs may stimulate “priming”, that is, enhancement of immune response upon second encounter, and (3) myeloid cells respond earlier to acute insults prior to utilization of lymphoid cells. Therefore, for the third part, we challenged bEV-primed young mice with *E. coli* antigens (LPS, TSST-1) and checked which myeloid populations are upregulated in response to this “second” encounter. tSNE maps are shown in Figure 5, in which we included the subset of normal environment mouse populations, both fetal and young “primed” mice. A summary of the population incidence changes across different setups is shown in Table 4 and 5.

**Figure 5.**
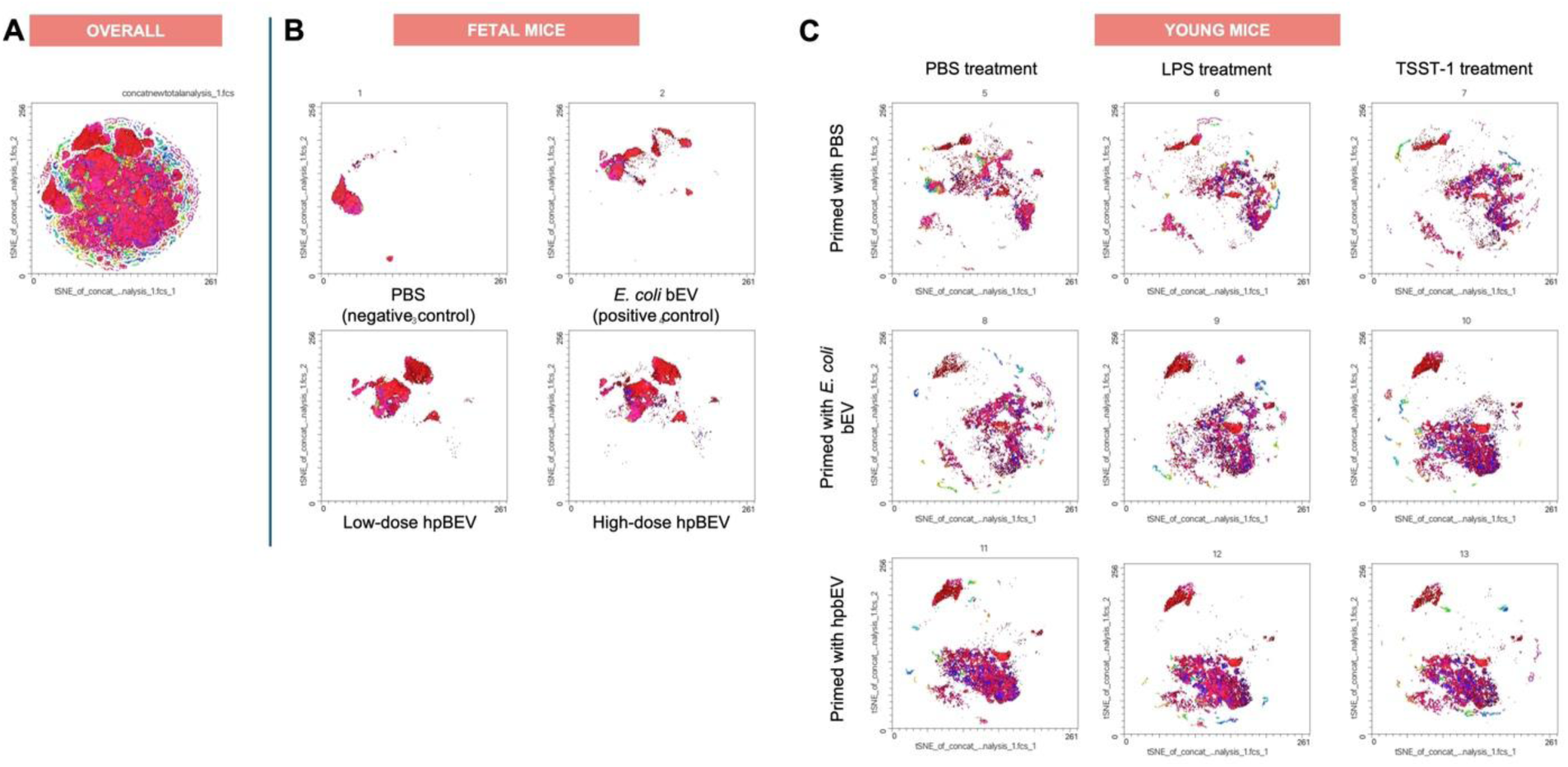
tSNE graphs for the myeloid population in the germ-free mice and the young mice setups. The overall tSNE map is shown (A), which is comprised of all cell populations identified from the germ-free mice setups (B) and the young mice setups (C). In the germ-free mice setups (B), we observe drastically different tSNEs across different treatment groups, although hpbEVs appear to activate similar populations. In the young mice setups (C), each priming agent, activates different populations entirely. Across different challenge treatments (LPS, TSST-1), cell populations remain apparently similar to controls (PBS), with the presence and presence of some populations. PBS, phosphate buffered saline. bEV, bacterial extracellular vesicles. hpbEV, human placental bacterial extracellular vesicles. LPS, lipopolysaccharide. TSST-1, toxic shock syndrome toxin-1.

**Table 4.** Cell clusters observed in the CyTOF analysis of myeloid cells in young, primed mice.

| CLUSTER | MYELOID MARKERS |
| --- | --- |
| 1 | CD11c c-kit XCR1 CD14 CD11b FceR1a CD8a Gr-1 CD86 |
| 2 | CD11c Arg1 c-kit XCR1 CD14 CD11b FceR1a CD193 CD8a Ly-6g Gr-1 CD103 CD86 CD206 F4-80 |
| 3 | c-kit XCR1 CD86 CD206 |
| 4 | CD11c Arg1 c-kit XCR1 FceR1a Gr-1 CD206 |
| 5 | Arg1 c-kit XCR1 CD14 CD11b FceR1a CD193 CD8a Ly-6g Gr-1 CD103 CD86 CD206 SiglecF F4-80 |
| 6 | CD11c XCR1 CD11b CD86 CD206 |
| 7 | CD11c Arg1 XCR1 CD86 CD206 |
| 8 | CD11c |
| 9 | CD11c XCR1 FceR1a CD86 CD206 |
| 10 | XCR1 CD86 CD206 F4-80 |
| 11 | XCR1 |
| 12 | CD11c XCR1 |
| 13 | CD11c c-kit XCR1 CD86 CD206 Siglec-F |
| 14 | XCR1 CD86 |
| 15 | CD11c c-kit XCR1 CD103 CD86 CD206 |
| 16 | CD11c Arg1 c-kit XCR1 CD11b FceR1a CD193 GR-1 CD86 CD206 |
| 17 | CD11c Arg1 CD11b CD86 CD206 |
| 18 | CD11c XCR1 CD11b CD86 F4-80 |
| 19 | CD11c XCR1 CD14 CD11b FceR1a CD193 CD8a Ly-6g Gr-1 CD103 CD86 CD206 Siglec-F F4-80 |
| 20 | CD11c Gr-1 CD86 CD206 |
| 21 | XCR1 MHC-II |
| 22 | -- |
| 23 | CD86 |
| 24 | CD11c Arg1 XCR1 CD11b FceR1a CD193 CD8a Ly-6g Gr-1 CD86 |
| 25 | CD11b |
| 26 | Arg1 CD11b FceR1a CD206 |
| 27 | CD11b Ly-6g Gr-1 CD86 |
| 28 | CD11c |
| 29 | CD11c Arg1 c-kit XCR1 CD14 CD11b FceR1a Gr-1 |
| 30 | CD11c Arg1 c-kit XCR1 CD14 CD11b FceR1a CD193 CD8a Ly-6g Gr-1 CD86<br>CD206 Siglec-F |

Heat maps for the myeloid population are shown in Figure 6. Overall, two-way ANOVA suggests that most of the clusters that significantly change in mean counts across various treatments occur in Clusters 1, 2, and 3 (p < 0.0001). Cluster 1 (c-kit+ XCR1+ CD14+ CD11c+ CD11b+ FceR1a+ CD8a+ Gr-1+ CD86+) and Cluster 2 (c-kit+ XCR1+ CD14+ CD11c+ CD11b+ FceR1a+ CD8a+ Gr-1+ CD86+ Arg1+ CD193+ Ly-6g+ CD103+ CD206+ F4-80+) both represent widely-marked myeloid progenitor cell populations. In both clusters, significant differences in the mean populations are observed against all young mouse myeloid cells across all treatments, strongly indicating that these cell types are only found in fetal environments and in embryonic tissues. Comparing across fetal populations, it is interesting to note that both populations are not significantly different in (1) control conditions *vs*. a high dose of hpbEVs (p > 0.9999) and (2) EC bEV treated mice *vs*. a low dose of hpbEVs (p > 0.9999).

**Figure 6.**
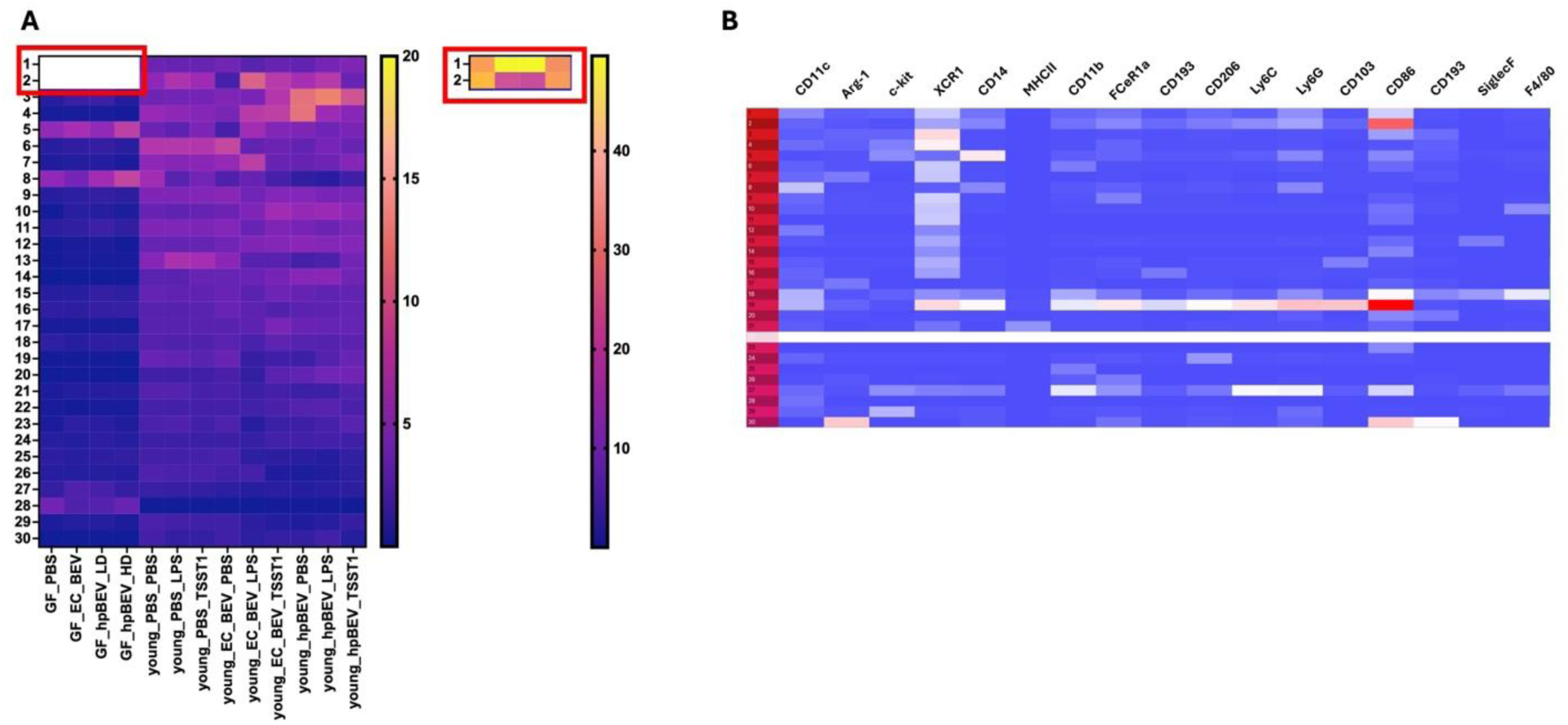
Heat maps for the myeloid populations in the germ-free mice setups and the young mice setups. (A) Each population can be broken down into different clusters, as shown in the heatmap (blue palette). Overall, due to the large number of populations (200), the study was limited only to the top 30 most abundant clusters for succeeding analyses. The relative abundances of the different clusters, or cell types, vary across different treatments, as shown in the heatmap (dark blue palette). Relatively, clusters 1 and 2 in the germ-free mice setups are significantly higher (two-way ANOVA, p <0.05) compared to most of the populations within their setups and across different treatments. Clusters 3-8 significantly differ in abundances (p < 0.05) across some of the treatments, distributed between germ-free and young mice setups. Clusters 9-30 do not vary significantly across all treatments (p > 0.05). (B) Each cluster represents a specific cell type, whose marker positivities vary across each marker. A summary of all markers is presented in Table 5 in the text. PBS, phosphate buffered saline. EC bEV, *E. coli* bacterial extracellular vesicles. hpbEV, human placental bacterial extracellular vesicles. LPS, lipopolysaccharide. TSST-1, toxic shock syndrome toxin-1.

**Table 5.** Comparisons across different setups of myeloid populations, and the absolute differences between the two populations shown using two-way ANOVA. A significant difference is denoted by p < 0.05.

| Setups | Comparator | Difference | Significance |
| --- | --- | --- | --- |
| **Cluster 3 (c-kit+ XCR1+ CD86+ CD206+)** |  |  |  |
| **Fetal (PBS)** | Young hpBEV-primed, PBS exposed | -10.91 | 0.0003 |
|  | Young hpBEV-primed, LPS exposed | -12.19 | <0.0001 |
|  | Young hpBEV-primed, TSST-1 exposed | -8.835 | 0.0103 |
| **Fetal (EC bEV)** | Young hpBEV-primed, PBS exposed | -10.29 | 0.0008 |
|  | Young hpBEV-primed, LPS exposed | -11.57 | <0.0001 |
|  | Young hpBEV-primed, TSST-1 exposed | -8.22 | 0.026 |
| **Fetal (LD hpbEV)** | Young hpBEV-primed, PBS exposed | -10.23 | <0.0001 |
|  | Young hpBEV-primed, LPS exposed | -11.5 | <0.0001 |
|  | Young hpBEV-primed, TSST-1 exposed | -8.153 | 0.0066 |
| **Fetal (HD hpbEV)** | Young hpBEV-primed, PBS exposed | -10.99 | <0.0001 |
|  | Young hpBEV-primed, LPS exposed | -12.27 | <0.0001 |
|  | Young hpBEV-primed, TSST-1 exposed | -8.918 | 0.0015 |
| Young PBS primed,<br>PBS exposed | Young hpBEV-primed, PBS exposed | -8.117 | 0.007 |
|  | Young hpBEV-primed, LPS exposed | -9.393 | 0.0006 |
| Young PBS primed,<br>LPS exposed | Young hpBEV-primed, PBS exposed | -7.93 | 0.0098 |
|  | Young hpBEV-primed, LPS exposed | -9.207 | 0.0008 |
| Young PBS primed,<br>TSST-1 exposed | Young hpBEV-primed, PBS exposed | -7.883 | 0.0106 |
|  | Young hpBEV-primed, LPS exposed | -9.16 | 0.0009 |
| Young EC-bEV primed,<br>PBS exposed | Young hpBEV-primed, PBS exposed | -7.357 | 0.0258 |
|  | Young hpBEV-primed, LPS exposed | -8.633 | 0.0026 |
| Young EC-bEV primed,<br>LPS exposed | Young hpBEV-primed, PBS exposed | -7.587 | 0.0177 |
|  | Young hpBEV-primed, LPS exposed | -8.863 | 0.0017 |
| Cluster 4 (CD44+) |  |  |  |
| Fetal (PBS) | Young hpBEV-primed, PBS exposed | -11.33 | 0.0001 |
| Fetal (EC bEV) | Young hpBEV-primed, PBS exposed | -11.32 | 0.0001 |
| Fetal (LD hpbEV) | Young EC bEV-primed, LPS exposed | -7.161 | 0.0351 |
|  | Young EC bEV-primed, TSST-1 exposed | -7.548 | 0.0189 |
|  | Young hpbEV-primed, PBS exposed | -11.29 | <0.0001 |
| Fetal (HD hpbEV) | Young EC bEV-primed, LPS exposed | -7.238 | 0.0312 |
|  | Young EC bEV-primed, TSST-1 exposed | -7.625 | 0.0166 |
|  | Young hpbEV-primed, PBS exposed | -11.37 | <0.0001 |
| Cluster 6 (CD69+ PD-1+ CD44+ CD3+ FOXP3+ CD38+) |  |  |  |
| Fetal (PBS) | Young EC bEV-primed, PBS exposed | -7.792 | 0.047 |
| Fetal (LD hpbEV) |  | -7.487 | 0.0209 |
| Fetal (HD hpbEV) |  | -7.737 | 0.0137 |
| Cluster 7 (CD69+ CD44+ CCR7+ CD24+) |  |  |  |
| Fetal (LD hpbEV) | Young EC bEV-primed, LPS exposed | -7.197 | 0.0332 |
| Fetal (HD hpbEV) |  | -7.412 | 0.0236 |
| Cluster 8 (CCR7+) |  |  |  |
| Fetal (HD hpbEV) | Young hp bEV-primed, PBS exposed | 7.433 | 0.0228 |
|  | Young hp bEV-primed, LPS exposed | 7.653 | 0.0158 |
|  | Young hp bEV-primed, TSST-1 exposed | 6.933 | 0.0496 |

In Cluster 3 (c-kit+ XCR1+ CD86+ CD206+), markers for monocytes/macrophages in the intestinal epithelia, there are significant differences between fetal and young mice. These cells are activated more in hpbEV-primed young mice compared to fetal mice, and only certain treatment groups in young mice have more of these cellpopulous. Interestingly, these cells do not significantly differ between fetal mouse setups.

Various dendritic cell groups, characterized by CD11c+ CD206+ and varying other markers, have varying population differences across different setups. Among treatment groups with significant differences, Clusters 4,6, and 7 generally showed lower counts in fetal setups versus young bEV-primed mice. Interestingly, Cluster 8, which is the only single-marked cluster (CD11c+ only), showed higher population counts in fetal setups compared to young hpbEV-primed mice.

In summary, our data suggest that upon second encounter, embryonic progenitors, macrophage/dendritic-cell-like cells, and dendritic cell-like populations dominate the myeloid response in GF mice. In contrast, in NE mice, a more targeted macrophage-like and dendritic cell-like response dominates.

## 4. Discussion

We established that bEVs influence fetal mucosal immunity and foster immune programming for a more targeted response to subsequent challenges. Introducing bEVs into the fetal gut drove distinct immune cell differentiation programs, as shown by CyTOF analysis in murine models. In conventionally colonized mice, bEV exposure led to increased memory and regulatory markers in lymphoid populations, with a concomitant dose-dependent expansion of monocyte/dendritic and neutrophil-like lineages in myeloid clusters. In GF setups, this exposure primarily activated granulocytic-like cells and shifted lymphocytes toward early memory phenotypes. Upon a second encounter with *E. coli* antigens, progenitor cells, macrophage/dendritic-like populations, and dendritic cell subsets dominated the myeloid response.

In the intestine, multiple lymphoid microenvironments exist, including the mucosa-associated lymphoid tissue (MALT), the lamina propria, and the epithelial cell tracts lined with intraepithelial lymphocytes (IEL)^61^. *Via in utero* fetal swallowing, any bEVs introduced in the amniotic sac can be transferred into the fetal gut and activate naïve immunity in the tissues. Our observations in lymphoid and myeloid cells led us to speculate that bEVs stimulate the immune environment in the murine fetal gut. Our findings have the potential to advance our understanding of how maternal inflammation during pregnancy programs long-term immune output from hematopoietic progenitors and subsequently impacts offspring immune function.

In the lymphoid population, we identified five CD45+ populations in the fetal gut of normal-environment mice. One interesting population is likely lymphoid progenitors (Cluster 22) expressing multiple markers of various canonical lymphocytes (CD45+ CD3+ CD8+ NK1.1+ CD25+ CD69+ PD-1+ CD44+ CD62L+ CCR7+ CXCR3+ GATA3+ CXCR4+ FoxP3+ CD24+CD27+ TCRyd+). The conventional outline for lymphocyte differentiation starts from the hematopoietic stem cells, differentiating into lymphoid-primed multipotent progenitors (LMPP), to common lymphoid progenitors (CLP), to committed progenitors, and finally to mature lymphocytes^62^. The slew of markers in this cluster may point towards a new lymphoid progenitor phenotype in the intestines that can readily differentiate into specific lineages, as needed. In our GF mice, we also observe a similar lymphoid cluster bearing multiple markers (Cluster 29). The relatively low incidence of these cells indicates they may represent transient populations since extrathymic development in the intestines is naturally repressed in the presence of a normal thymus^63^.

PD-1 is a marker typically associated with exhaustion, but may alternatively represent a marker of activation and chronic stimulation^64,65^. Both clusters decrease in incidence with increasing bEV exposure, possibly indicating a decrease in exhausted phenotypes with concomitant release from PD-1 inactivation that would allow for enhanced action in response to bEV exposure. We also observe a small population of double-positive T cells (CD45+ CD4+ CD8+) in both NE and GF mice (Cluster 33 and Cluster 28, respectively) that increases with exposure to increasing doses of bEVs. Albeit relatively low in incidence, we postulate that double-positive T cells may play important roles in fetal immunity due to their memory nature as well as their potential to produce higher levels of cytokines and chemokines compared to CD4+ or CD8+ only cells^66,67^.

In the intestine, CD24+ CD44+ cells represent fetal intestinal stem cells (ISCs), which are important in potentiating the self-renewing capacity of the tissue to respond to antigenic challenges and injury. Although Lgr5 is a more putative marker for ISCs, CD24 provides an acceptable marker for identifying stem cells in mice, with similar counterparts in human intestinal cells ^68–71^. ISCs provide crosstalk with other lymphocytes for further differentiation into Paneth cells or tuft cells. Paneth cells are secretory cells that may aid intestinal defense by secreting granules containing a host of defensive proteins, including antimicrobial peptides and enzymes^72–74^. ISCs can also express MHCs, allowing them to serve as adjunct antigen-presenting cells for T-cell activation and proliferation^75,76^. We observed a general increase in the incidence of the clusters upon bEV exposure compared to negative controls, suggesting that bEV presence preemptively induces preparation of the intestines for possible damage. However, the presence of CD45 antigen in the CD24+ cluster strongly suggests that this is a lymphocyte population. A population of TCRɣ*δ* cells has been demonstrated to produce a similar phenotype, but the absence of a TCRɣ*δ* marker in this population suggests otherwise^77^. It might be that the populations are an early phenotype of naïve T cells that do not express CD4+ or CD8+ yet, but are early activated via CD44 upregulation during encounter with antigens^78^.

As for the CD45- cells, the majority of the populations express CD44 and CCR7, in combination with other markers; interestingly, the populations would either have only these markers present or have them in combination with a few markers that would be incomplete for us to classify them into one of the canonical lymphocyte types. In line with the above hypothesis, we suspect that the majority of the clusters are transition states into other lymphocytes with early expression of CD44, CCR7, and/or CD62L as early memory or activation markers.

In general, we observe the predominant upregulation of markers as bEV exposure increases, displaying the capacity of bEVs to induce memory and/or activation phenotypes in these cell clusters. Therefore, two populations of naïve memory populations are fine-tuned by CD45 expression: (1) CD45+ populations are cells trending towards maturity *via* increasing survival signal receptivity, while (2) CD45-populations are cells that are geared toward an augmented inflammatory response to the presence of bEVs^79–82^. However, in our GF mice, we do not see any significant differences between negative controls and bEV-exposed pups in the CD45+ CD24+ CD44+ cluster, instead we see a similar trend in the increase with the CD45-CD24+ CD44+ cluster. Therefore, if we reduce any influences from the environment, the acute course of fetal murine gut immunity is to expand inflammatory responses against the presence of bEVs, while maintaining all populations in the course of maturation.

In the myeloid population, we observe six (6) clusters of CD11b+ cells in the NE mice. The choice in utilizing CD11b as a primary gate is secondary to the abundance of CD11b-expressing monocytes, macrophages, and dendritic cells that lie within the murine epithelium at the start of myeloid-derived cell residency coming from the fetal liver and blood^83–85^. Three clusters (Cluster 3, 12, 21) display multiple markers reminiscent of the lymphoid progenitor population we have postulated previously. The observed markers correspond to dendritic cells (CD11b+, CD103+) and macrophages/monocytes (F4/80+, CD14+, CD86+, XCR1+, Ly6C+, Gr-1+), with a few uncommon markers as well (CD8+, FcεRIα+, CD193+, Siglec-F+)^83–89^. We purport that these cells may be monocyte-dendritic progenitors, which can produce monocyte-derived dendritic cells^90,91^. Interestingly, we do not see a strong expression of MHCII in any of the cell clusters, potentially indicating that the levels of bEV are not high enough to induce antigen presentation, and that development of exposed immune cells in the gut may be instead driven by direct interactions with bEVs; this remain to be proven *via* functional studies. Another alternative explanation is that MHCII is in the myeloid panel where antibodies against epithelial cells (CD24, CD44) were not accounted for; indeed, Paneth cells and crypt villi in the intestine natively express MHCII and may play important roles in antigen presentation^92,93^; future research may also attempt to elucidate the role of these cells in inflammation and immune education upon bEV exposure. Nonetheless, we see a similar progenitor cluster in GF mice (Cluster 7) that also observes the same trend, indicating that this population may be an important player in myeloid immunity.

There are three clusters of cells (Myeloid Cluster 1, 16, 19) that are upregulated in low concentrations of bEVs but downregulated upon exposure to high concentrations in NE mice. Although closely resembling some markers present in the previously mentioned monocyte-dendritic progenitors, the fewer markers present in the clusters, as well as their differential expression, may point to an entirely different population. We suspect that the clusters may represent intestinal macrophages (CD11b+/− F4/80+ CD86+ CD11c+ XCR1+). It is yet unclear why there is a concentration-dependent decrease in the incidence of these cells. Still, it can be speculated that there might be a switch towards a different phenotype of macrophages that were not captured by our panel. Possible alternative populations may include (1) CD169+ macrophages, which are mainly regulatory in nature^94,95^, and (2) TCR+ macrophages, which are TNF-tuned, CCL2-releasing, and highly phagocytic subpopulations^96,97^. We observe a similar macrophage cluster in RG mice (Cluster 14) that also exhibits the same trend.

Interestingly, we see an increase in neutrophils (Ly6G+) in both groups of mice (Cluster 11 for NE mice, Clusters 16, 19, 25 in GF mice). Upon acute inflammation, resident monocytes in the intestine recruit neutrophils to aid in the response against pathogens^98,99^. Neutrophils can release chemokines that can recruit a second wave of macrophages^100^; however, as mentioned previously, neutrophil-assisted recruitment of macrophages may only contribute to an increase of macrophage counts in lower concentrations of bEVs.

For the GF mice, an interesting cluster is the B-cell-like population, characterized by CD19⁺ CD24⁺. This mimics a tolerogenic phenotype, similar to an IL-10 suppressed Treg cell; in an experimental model (collagen-induced arthritis), the adoptive transfer of CD19⁺CD24⁺ transitional B cells reduced disease severity via secretion of IL-10 in an inflammatory environment^101^. It is attractive to think that in an environment without native influences, Bregs contribute to maintenance of homeostatic neutrality that prevents over-exertion of the immune system to a transient exposure and may serve to also train the immune system ^102,103^.

Overall, we observe differentially expressed populations in the pups of controls versus bEV treated mice, with general upregulation of a majority populations as bEV concentration increases. Lymphoid progenitors, intestinal stem cells, and monocyte-dendritic progenitors are increased in both NE and GF mice, indicating that bEV exposure has the propensity to produce a pool of cells readily available for targeted differentiation and maturation. Exhausted phenotypes and memory/activated populations also decrease upon bEV exposure. Macrophages are produced in a concentration-dependent manner, with lower concentrations of bEVs providing higher incidences compared to higher concentrations. Neutrophil production is also stimulated upon bEV exposure, but it remains to be seen whether cells are mature enough to contribute to fetal intestinal immunity.

Our experiments with young mice provide insights as to how prenatal bEV stimulation may modulate second encounter responses. Similar to what we have seen in myeloid populations in embryonic conditions, a majority of the populations upregulated in young mice are progenitors characterized by multiple cellular markers (Cluster 1 and Cluster 2). There are two important points with these progenitors: (1) they are highly present in fetal life at baseline, as indicated by the absence of significant differences among embryonic controls, and (2) they do not persist in later life, as seen with the significant decreases from fetal setups to young mice setups. We can potentially think of these cells as native progenitors resting within the murine intestine that slowly differentiate into other cells over time. Whether the progenitors function differently compared to the aforementioned CD45+ intestinal stem cells is yet unknown. In both clusters, there are two interesting scenarios where we see no difference in progenitor counts. In high-dose hpbEVs versus controls, we theorize that the high dose leads to an immune response that decreases the antigen burden and decreases the recruitment of progenitors. In mice treated with EC bEV versus a low dose of hpbEV, we theorize that both exposures are biological equivalents of each other leading to no-difference in progenitor counts upon encounter.

Cluster 3 may represent dendritic cells in the intestine that are activated upon introduction of an unrecognized antigen. CD86 and CD206 are canonical macrophage markers for M1 and M2 macrophages, respectively ^104^. However, c-kit positivity is an essential feature of dendritic cells ^105^ The presence of both CD86 and CD206 markers points more toward a monocyte-derived dendritic cell. Noticeably, only hpbEV-primed setups show a significant increase in population counts regardless of exposure status in young life. Therefore, we postulate that in response to hpbEV exposure, dendritic cells become activated to serve as an initial defense mechanism for the mucosal environment. However, the lack of MHC II positivity in these cells may imply the beginning of clonal expansion upon initial contact.

Other dendritic cell-like phenotypes also increase upon bEV exposure. In CD44+ clusters, we see that hpbEV primed cells have baseline higher CD44+ cells compared to fetal setups. ECbEV-primed setups have notably higher CD44+ counts compared to hpbEV-exposed fetal mice. We can speculate that the CD44+ cells are higher in primed mice due to their relatively longer exposure time with the hpbEVs compared to embryonic ones. Interestingly, EC bEV-primed mice also showed higher CD44+ counts versus hpbEV exposed fetal mice, showing that subsequent exposure to LPS/TSST-1 in EC bEV-primed mice may trigger activation of a primed state that enables a higher response to a secondary exposure.

Cluster 6 may also represent progenitor cells, as the cluster expresses neutrophil cell (CD38+) and dendritic cell markers (CD69+ CD44+). CD3+ expression is a marker of macrophage cells^106^ while regulation of FOXP3 in myeloid cells is necessary for differentiation and maturation. Deletion of PD-1 in myeloid cells stimulates an increase in T effector memory cell proliferation, increasing memory responses among T cells with improved functionality^107^. In tumor infiltrating myeloid cells, PD-1 positively regulates the differentiation of myeloid-derived suppressive cells, subsequently leading to a suppression of T effector cells population and function. The expression of this cluster increases in EC-bEV primed, PBS exposed mice, implying that EC-bEV priming, on baseline, stimulates an increase in suppressive clusters. Non-difference upon priming with hpbEV and LPS/TSST post-exposure in EC bEV young mice may imply that (1) hpbEV exposure does not stimulate production of suppressive clusters and (2) LPS/TSST post-exposure results in dampening of suppressors to promote an increased response upon recognition of these antigens.

CCR7+ is an inflammatory marker expressed by myeloid dendritic cells and lymphoid cells, resulting in chemotaxis of recruited cells ^108,109^. Thus, Cluster 8 cells may represent myeloid cells that are increased secondary to a controlled inflammation from hpbEV priming. It is attractive to think that the decrease in hpbEV-primed setups in Cluster 1 and 2 is due to differentiation to these monocyte-dendritic cell hybrids; however, further trajectory analysis would be required to confirm this hypothesis. Interestingly, the observed increases are mild and may thus represent an early inflammatory response to priming and/or antigen exposure.

Notably different from the primed setups *versus* embryonic fetal setups is the absence of purely macrophage or neutrophil populations. In fetal life, fetal macrophages, neutrophils, and monocytes can respond to antigens as the major pro-inflammatory cells in the developing fetus^89,110^. We identified several clusters that may be macrophage and neutrophil candidates, although the clusters do not vary significantly between different setups. We speculate that these cells may be more apparent in later time points, since it is expected that dendritic cells would be the first responders to a foreign antigen, subsequently activating other myeloid cells to initiate an immune response^111,112^. It may also be advantageous that the differentiated cells are kept to a lower state of activation of proliferation to prevent unwarranted, widespread immune response within the gut and promote a constant low exposure of antigens that can be exploited for intestinal immune education^112^.

Our study has certain limitations. First, it is notable that most of our populations do not readily fit into the classical descriptions of conventional immune cells in terms of surface markers. Although we provide the phenotypic landscape of the immune repository upon exposure to bacterial extracellular vesicles, functional studies are still necessary to provide context as to how these cells may act locally in response to a relatively mild immunogen. Second, we also utilized mice as our subjects due to their convenience and manipulability. Although this model may explain how humans would also react toward bEV exposure, studies utilizing human-derived cells or a humanized mouse model may be more representative. Third, interaction with the environment will also play a crucial role in modifying the early immunophenotype of our models. Therefore, studies that employ purely GF mice under GF environments will provide a more conclusive look into how naïve immunity is affected by environmental changes and the evolving microbiome. Fourth, our approach also utilizes a simple k-means clustering network – more powerful approaches toward dissecting the overall population may be warranted in future studies, and trajectory analysis will determine how one phenotype of cells evolves into other phenotypes. Finally, additional research, especially functional studies, are needed to further elucidate the exact mechanisms of fetal immune development upon bEV exposure.

As the establishment of the presence of placental bEVs has only come into recent light, this study addresses the lack of studies into the importance of bEVs on the developing immune system. Our study provides an initial look at how bEVs can influence protective responses in early life. In summary, we see that (1) progenitors are highly prominent during fetal life, (2) there is a decrease of progenitor populaitnos upon birth going to young life, and (3) there is a propensity for myeloid cells to differentiate into antigen-presenting cells in a primed response – all of which are expected trends in immune system development upon exposure to an antigen, such as bEVs. This study also raises several important research avenues to address in the future, including (1) Which specific populations do the identified cell populations differentiate into upon longer exposure? (2) What are the corresponding lymphoid responses in the primed setup? (3) Would responses between animal and human immunity be the same? (4) What are the functional responses of an EV-primed system against other antigens? and (5) What is the value of bEVs and any other modifications in effecting a memory response similar to a vaccine? The last point is a relevant point, since EVs can provide a more sustainable and convenient approach to vaccination^113,114^Finally, an exciting question is whether EVs coming from various sources (plants, viruses, bacteria) would provoke similar or discrete responses in early-life immunity and whether the responses can be harnessed to develop more targeted immune tolerance/immunity against specific pathogens.

## 5. Conclusion

This study elucidates the role of bEVs in shaping fetal immune development. We observed key immune trends, including the prominence of progenitor cells during fetal life, their decline postnatally, and the primed differentiation of myeloid cells into antigen-presenting cells. Our findings align with expected immune maturation patterns following antigenic exposure. Future studies should explore the long-term differentiation of these cells, their interaction with lymphoid responses, and the potential for bEVs to induce memory-like immunity. Given their stability and bioavailability, bEVs may serve as a novel platform for immune modulation, with applications in neonatal immunity, vaccine development, and immune tolerance strategies.

## Acknowledgment

We sincerely thank Ms. **Talar Kechichian** for her invaluable assistance in securing the animal amendment in a timely manner. Ms. **Phyllis Gamble** for her dedication to maintaining the animal colony. Additionally, we are grateful to Dr. **Enkhtuya Radnaa, Ph.D.,** for her expertise and support in performing surgical procedures. The contributions of the aforelisted individuals were instrumental in the successful completion of this study.

## Conflict of interest

Authors declare there is no conflict of interest.

## Data Availability Statement

Mass Cytometry Data are available in https://doi.org/10.5281/zenodo.15658656 These datasets provide comprehensive insights into the cellular composition and protein expression profiles analyzed through mass cytometry techniques.

## Consent of publication

Not applicable

## Funding

The Robert and Janice McNair Foundation, R01 HD109095, and R01 HD109780 to S.A.B. R01HD100729-05 to RM

## Notes

### Competing Interest Statement

The authors have declared no competing interest.

## References

1. Stout MJ, Conlon B, Landeau M, et al. Identification of intracellular bacteria in the basal plate of the human placenta in term and preterm gestations. Am J Obstet Gynecol. 2013;208(3):226 e221–227.

2. PrabhuDas M, Bonney E, Caron K, et al. Immune mechanisms at the maternal-fetal interface: perspectives and challenges. Nat Immunol. 2015;16(4):328–334.

3. Liu S, Diao L, Huang C, Li Y, Zeng Y, Kwak-Kim JYH. The role of decidual immune cells on human pregnancy. J Reprod Immunol. 2017;124:44–53.

4. Mor G, Kwon JY. Trophoblast-microbiome interaction: a new paradigm on immune regulation. Am J Obstet Gynecol. 2015;213(4 Suppl):S131–137.

5. Lash GE. Molecular Cross-Talk at the Feto-Maternal Interface. Cold Spring Harb Perspect Med. 2015;5(12).

6. Gomez-Lopez N, StLouis D, Lehr MA, Sanchez-Rodriguez EN, Arenas-Hernandez M. Immune cells in term and preterm labor. Cell Mol Immunol. 2014;11(6):571–581.

7. Alijotas-Reig J, Llurba E, Gris JM. Potentiating maternal immune tolerance in pregnancy: a new challenging role for regulatory T cells. Placenta. 2014;35(4):241–248.

8. Nancy P, Tagliani E, Tay CS, Asp P, Levy DE, Erlebacher A. Chemokine gene silencing in decidual stromal cells limits T cell access to the maternal-fetal interface. Science. 2012;336(6086):1317–1321.

9. Williams PJ, Searle RF, Robson SC, Innes BA, Bulmer JN. Decidual leucocyte populations in early to late gestation normal human pregnancy. J Reprod Immunol. 2009;82(1):24–31.

10. Harris LK, Benagiano M, D’Elios MM, Brosens I, Benagiano G. Placental bed research: II. Functional and immunological investigations of the placental bed. Am J Obstet Gynecol. 2019;221(5):457–469.

11. Rouas-Freiss N, Moreau P, LeMaoult J, Papp B, Tronik-Le Roux D, Carosella ED. Role of the HLA-G immune checkpoint molecule in pregnancy. Hum Immunol. 2021;82(5):353–361.

12. Jacobs SO, Sheller-Miller S, Richardson LS, Urrabaz-Garza R, Radnaa E, Menon R. Characterizing the immune cell population in the human fetal membrane. Am J Reprod Immunol. 2021;85(5):e13368.

13. True H, Blanton M, Sureshchandra S, Messaoudi I. Monocytes and macrophages in pregnancy: The good, the bad, and the ugly. Immunol Rev. 2022;308(1):77–92.

14. Motomura K, Hara M, Ito I, Morita H, Matsumoto K. Roles of human trophoblasts’ pattern recognition receptors in host defense and pregnancy complications. J Reprod Immunol. 2023;156:103811.

15. Muralidhara P, Sood V, Vinayak Ashok V, Bansal K. Pregnancy and Tumour: The Parallels and Differences in Regulatory T Cells. Front Immunol. 2022;13:866937.

16. Xie M, Li Y, Meng YZ, et al. Uterine Natural Killer Cells: A Rising Star in Human Pregnancy Regulation. Front Immunol. 2022;13:918550.

17. Levy M, Blacher E, Elinav E. Microbiome, metabolites and host immunity. Curr Opin Microbiol. 2017;35:8–15.

18. Netea MG, Joosten LA, Latz E, et al. Trained immunity: A program of innate immune memory in health and disease. Science. 2016;352(6284):aaf1098.

19. Mandal M, Donnelly R, Elkabes S, et al. Maternal immune stimulation during pregnancy shapes the immunological phenotype of offspring. Brain Behav Immun. 2013;33:33–45.

20. Mishra A, Lai GC, Yao LJ, et al. Microbial exposure during early human development primes fetal immune cells. Cell. 2021;184(13):3394–3409 e3320.

21. Zengeler KE, Lukens JR. Innate immunity at the crossroads of healthy brain maturation and neurodevelopmental disorders. Nat Rev Immunol. 2021;21(7):454–468.

22. Jash S, Sharma S. In utero immune programming of autism spectrum disorder (ASD). Hum Immunol. 2021;82(5):379–384.

23. Scott RL, Vu HTH, Jain A, Iqbal K, Tuteja G, Soares MJ. Conservation at the uterine-placental interface. Proc Natl Acad Sci U S A. 2022;119(41):e2210633119.

24. Hodyl NA, Stark MJ, Osei-Kumah A, Clifton VL. Prenatal programming of the innate immune response following in utero exposure to inflammation: a sexually dimorphic process? Expert Rev Clin Immunol. 2011;7(5):579–592.

25. Langley-Evans SC, Carrington LJ. Diet and the developing immune system. Lupus. 2006;15(11):746–752.

26. Lopez DA, Apostol AC, Lebish EJ, et al. Prenatal inflammation perturbs murine fetal hematopoietic development and causes persistent changes to postnatal immunity. Cell Rep. 2022;41(8):111677.

27. Zhang Y, Bi J, Huang J, Tang Y, Du S, Li P. Exosome: A Review of Its Classification, Isolation Techniques, Storage, Diagnostic and Targeted Therapy Applications. Int J Nanomedicine. 2020;15:6917–6934.

28. Chen YF, Luh F, Ho YS, Yen Y. Exosomes: a review of biologic function, diagnostic and targeted therapy applications, and clinical trials. J Biomed Sci. 2024;31(1):67.

29. Shepherd MC, Radnaa E, Tantengco OA, et al. Extracellular vesicles from maternal uterine cells exposed to risk factors cause fetal inflammatory response. Cell Commun Signal. 2021;19(1):100.

30. Sheller-Miller S, Trivedi J, Yellon SM, Menon R. Exosomes Cause Preterm Birth in Mice: Evidence for Paracrine Signaling in Pregnancy. Sci Rep. 2019;9(1):608.

31. Menon R, Khanipov K, Radnaa E, et al. Amplification of microbial DNA from bacterial extracellular vesicles from human placenta. Front Microbiol. 2023;14:1213234.

32. Baglio SR, van Eijndhoven MA, Koppers-Lalic D, et al. Sensing of latent EBV infection through exosomal transfer of 5’pppRNA. Proc Natl Acad Sci U S A. 2016;113(5):E587–596.

33. Nabet BY, Qiu Y, Shabason JE, et al. Exosome RNA Unshielding Couples Stromal Activation to Pattern Recognition Receptor Signaling in Cancer. Cell. 2017;170(2):352–366 e313.

34. Torralba D, Baixauli F, Villarroya-Beltri C, et al. Priming of dendritic cells by DNA-containing extracellular vesicles from activated T cells through antigen-driven contacts. Nat Commun. 2018;9(1):2658.

35. Diamond JM, Vanpouille-Box C, Spada S, et al. Exosomes Shuttle TREX1-Sensitive IFN-Stimulatory dsDNA from Irradiated Cancer Cells to DCs. Cancer Immunol Res. 2018;6(8):910–920.

36. Fabbri M, Paone A, Calore F, et al. MicroRNAs bind to Toll-like receptors to induce prometastatic inflammatory response. Proc Natl Acad Sci U S A. 2012;109(31):E2110–2116.

37. Li X, Lei Y, Wu M, Li N. Regulation of Macrophage Activation and Polarization by HCC-Derived Exosomal lncRNA TUC339. Int J Mol Sci. 2018;19(10).

38. Liu J, Fan L, Yu H, et al. Endoplasmic Reticulum Stress Causes Liver Cancer Cells to Release Exosomal miR-23a-3p and Up-regulate Programmed Death Ligand 1 Expression in Macrophages. Hepatology. 2019;70(1):241–258.

39. Zhao S, Mi Y, Guan B, et al. Tumor-derived exosomal miR-934 induces macrophage M2 polarization to promote liver metastasis of colorectal cancer. J Hematol Oncol. 2020;13(1):156.

40. Loh JT, Zhang B, Teo JKH, et al. Mechanism for the attenuation of neutrophil and complement hyperactivity by MSC exosomes. Cytotherapy. 2022;24(7):711–719.

41. Thery C, Duban L, Segura E, Veron P, Lantz O, Amigorena S. Indirect activation of naive CD4+ T cells by dendritic cell-derived exosomes. Nat Immunol. 2002;3(12):1156–1162.

42. Hong X, Schouest B, Xu H. Effects of exosome on the activation of CD4+ T cells in rhesus macaques: a potential application for HIV latency reactivation. Sci Rep. 2017;7(1):15611.

43. Maybruck BT, Pfannenstiel LW, Diaz-Montero M, Gastman BR. Tumor-derived exosomes induce CD8(+) T cell suppressors. J Immunother Cancer. 2017;5(1):65.

44. Sheller-Miller S, Choi K, Choi C, Menon R. Cyclic-recombinase-reporter mouse model to determine exosome communication and function during pregnancy. Am J Obstet Gynecol. 2019;221(5):502 e501–502 e512.

45. Vidal MS, Jr., Radnaa E, Vora N, et al. Establishment and comparison of human term placenta-derived trophoblast cellsdagger. Biol Reprod. 2024;110(5):950–970.

46. Timmons BC, Reese J, Socrate S, et al. Prostaglandins are essential for cervical ripening in LPS-mediated preterm birth but not term or antiprogestin-driven preterm ripening. Endocrinology. 2014;155(1):287–298.

47. Baaten BJ, Tinoco R, Chen AT, Bradley LM. Regulation of Antigen-Experienced T Cells: Lessons from the Quintessential Memory Marker CD44. Front Immunol. 2012;3:23.

48. Campbell JJ, Murphy KE, Kunkel EJ, et al. CCR7 expression and memory T cell diversity in humans. J Immunol. 2001;166(2):877–884.

49. Kueberuwa G, Gornall H, Alcantar-Orozco EM, et al. CCR7(+) selected gene-modified T cells maintain a central memory phenotype and display enhanced persistence in peripheral blood in vivo. J Immunother Cancer. 2017;5:14.

50. Mullen KM, Gocke AR, Allie R, et al. Expression of CCR7 and CD45RA in CD4+ and CD8+ subsets in cerebrospinal fluid of 134 patients with inflammatory and non-inflammatory neurological diseases. J Neuroimmunol. 2012;249(1-2):86–92.

51. Unsoeld H, Pircher H. Complex memory T-cell phenotypes revealed by coexpression of CD62L and CCR7. J Virol. 2005;79(7):4510–4513.

52. Kohn LA, Hao QL, Sasidharan R, et al. Lymphoid priming in human bone marrow begins before expression of CD10 with upregulation of L-selectin. Nat Immunol. 2012;13(10):963–971.

53. Ivetic A, Hoskins Green HL, Hart SJ. L-selectin: A Major Regulator of Leukocyte Adhesion, Migration and Signaling. Front Immunol. 2019;10:1068.

54. Guiu J, Hannezo E, Yui S, et al. Tracing the origin of adult intestinal stem cells. Nature. 2019;570(7759):107–111.

55. Hou Q, Huang J, Ayansola H, Masatoshi H, Zhang B. Intestinal Stem Cells and Immune Cell Relationships: Potential Therapeutic Targets for Inflammatory Bowel Diseases. Front Immunol. 2020;11:623691.

56. Blasius AL, Barchet W, Cella M, Colonna M. Development and function of murine B220+CD11c+NK1.1+ cells identify them as a subset of NK cells. J Exp Med. 2007;204(11):2561–2568.

57. Wendel M, Galani IE, Suri-Payer E, Cerwenka A. Natural killer cell accumulation in tumors is dependent on IFN-gamma and CXCR3 ligands. Cancer Res. 2008;68(20):8437–8445.

58. Marquardt N, Wilk E, Pokoyski C, Schmidt RE, Jacobs R. Murine CXCR3+CD27bright NK cells resemble the human CD56bright NK-cell population. Eur J Immunol. 2010;40(5):1428–1439.

59. Yu L, Yang F, Zhang F, et al. CD69 enhances immunosuppressive function of regulatory T-cells and attenuates colitis by prompting IL-10 production. Cell Death Dis. 2018;9(9):905.

60. Cortes JR, Sanchez-Diaz R, Bovolenta ER, et al. Maintenance of immune tolerance by Foxp3+ regulatory T cells requires CD69 expression. J Autoimmun. 2014;55:51–62.

61. Gibbons DL, Spencer J. Mouse and human intestinal immunity: same ballpark, different players; different rules, same score. Mucosal Immunol. 2011;4(2):148–157.

62. Ghaedi M, Steer CA, Martinez-Gonzalez I, Halim TYF, Abraham N, Takei F. Common-Lymphoid-Progenitor-Independent Pathways of Innate and T Lymphocyte Development. Cell Rep. 2016;15(3):471–480.

63. Guy-Grand D, Azogui O, Celli S, et al. Extrathymic T cell lymphopoiesis: ontogeny and contribution to gut intraepithelial lymphocytes in athymic and euthymic mice. J Exp Med. 2003;197(3):333–341.

64. Sharpe AH, Pauken KE. The diverse functions of the PD1 inhibitory pathway. Nat Rev Immunol. 2018;18(3):153–167.

65. Jubel JM, Barbati ZR, Burger C, Wirtz DC, Schildberg FA. The Role of PD-1 in Acute and Chronic Infection. Front Immunol. 2020;11:487.

66. Pahar B, Lackner AA, Veazey RS. Intestinal double-positive CD4+CD8+ T cells are highly activated memory cells with an increased capacity to produce cytokines. Eur J Immunol. 2006;36(3):583–592.

67. Overgaard NH, Jung JW, Steptoe RJ, Wells JW. CD4+/CD8+ double-positive T cells: more than just a developmental stage? J Leukoc Biol. 2015;97(1):31–38.

68. Nefzger CM, Jarde T, Rossello FJ, et al. A Versatile Strategy for Isolating a Highly Enriched Population of Intestinal Stem Cells. Stem Cell Reports. 2016;6(3):321–329.

69. Gracz AD, Fuller MK, Wang F, et al. Brief report: CD24 and CD44 mark human intestinal epithelial cell populations with characteristics of active and facultative stem cells. Stem Cells. 2013;31(9):2024–2030.

70. Wang F, Scoville D, He XC, et al. Isolation and characterization of intestinal stem cells based on surface marker combinations and colony-formation assay. Gastroenterology. 2013;145(2):383–395 e381-321.

71. King JB, von Furstenberg RJ, Smith BJ, McNaughton KK, Galanko JA, Henning SJ. CD24 can be used to isolate Lgr5+ putative colonic epithelial stem cells in mice. Am J Physiol Gastrointest Liver Physiol. 2012;303(4):G443–452.

72. Lueschow SR, McElroy SJ. The Paneth Cell: The Curator and Defender of the Immature Small Intestine. Front Immunol. 2020;11:587.

73. Ouellette AJ. Paneth cells and innate mucosal immunity. Curr Opin Gastroenterol. 2010;26(6):547–553.

74. Satoh Y. Effect of live and heat-killed bacteria on the secretory activity of Paneth cells in germ-free mice. Cell Tissue Res. 1988;251(1):87–93.

75. Visan I. Stem cell-immune cell cross-talk. Nat Immunol. 2019;20(1):1.

76. Biton M, Haber AL, Rogel N, et al. T Helper Cell Cytokines Modulate Intestinal Stem Cell Renewal and Differentiation. Cell. 2018;175(5):1307–1320 e1322.

77. Sumaria N, Grandjean CL, Silva-Santos B, Pennington DJ. Strong TCRgammadelta Signaling Prohibits Thymic Development of IL-17A-Secreting gammadelta T Cells. Cell Rep. 2017;19(12):2469–2476.

78. Goldrath AW, Bogatzki LY, Bevan MJ. Naive T cells transiently acquire a memory-like phenotype during homeostasis-driven proliferation. J Exp Med. 2000;192(4):557–564.

79. Cho JH, Kim HO, Ju YJ, et al. CD45-mediated control of TCR tuning in naive and memory CD8(+) T cells. Nat Commun. 2016;7:13373.

80. Zikherman J, Doan K, Parameswaran R, Raschke W, Weiss A. Quantitative differences in CD45 expression unmask functions for CD45 in B-cell development, tolerance, and survival. Proc Natl Acad Sci U S A. 2012;109(1):E3–12.

81. McNeill L, Salmond RJ, Cooper JC, et al. The differential regulation of Lck kinase phosphorylation sites by CD45 is critical for T cell receptor signaling responses. Immunity. 2007;27(3):425–437.

82. Trowbridge IS, Thomas ML. CD45: an emerging role as a protein tyrosine phosphatase required for lymphocyte activation and development. Annu Rev Immunol. 1994;12:85–116.

83. Gross M, Salame TM, Jung S. Guardians of the Gut - Murine Intestinal Macrophages and Dendritic Cells. Front Immunol. 2015;6:254.

84. Bain CC, Bravo-Blas A, Scott CL, et al. Constant replenishment from circulating monocytes maintains the macrophage pool in the intestine of adult mice. Nat Immunol. 2014;15(10):929–937.

85. Farache J, Zigmond E, Shakhar G, Jung S. Contributions of dendritic cells and macrophages to intestinal homeostasis and immune defense. Immunol Cell Biol. 2013;91(3):232–239.

86. Bain CC, Scott CL, Uronen-Hansson H, et al. Resident and pro-inflammatory macrophages in the colon represent alternative context-dependent fates of the same Ly6Chi monocyte precursors. Mucosal Immunol. 2013;6(3):498–510.

87. Dunay IR, Damatta RA, Fux B, et al. Gr1(+) inflammatory monocytes are required for mucosal resistance to the pathogen Toxoplasma gondii. Immunity. 2008;29(2):306–317.

88. Shin JS, Greer AM. The role of FcepsilonRI expressed in dendritic cells and monocytes. Cell Mol Life Sci. 2015;72(12):2349–2360.

89. Lakhdari O, Yamamura A, Hernandez GE, et al. Differential Immune Activation in Fetal Macrophage Populations. Sci Rep. 2019;9(1):7677.

90. Yanez A, Coetzee SG, Olsson A, et al. Granulocyte-Monocyte Progenitors and Monocyte-Dendritic Cell Progenitors Independently Produce Functionally Distinct Monocytes. Immunity. 2017;47(5):890–902 e894.

91. Merad M, Sathe P, Helft J, Miller J, Mortha A. The dendritic cell lineage: ontogeny and function of dendritic cells and their subsets in the steady state and the inflamed setting. Annu Rev Immunol. 2013;31:563–604.

92. Wosen JE, Mukhopadhyay D, Macaubas C, Mellins ED. Epithelial MHC Class II Expression and Its Role in Antigen Presentation in the Gastrointestinal and Respiratory Tracts. Front Immunol. 2018;9:2144.

93. Jamwal DR, Laubitz D, Harrison CA, et al. Intestinal Epithelial Expression of MHCII Determines Severity of Chemical, T-Cell-Induced, and Infectious Colitis in Mice. Gastroenterology. 2020;159(4):1342–1356 e1346.

94. Oetke C, Vinson MC, Jones C, Crocker PR. Sialoadhesin-deficient mice exhibit subtle changes in B- and T-cell populations and reduced immunoglobulin M levels. Mol Cell Biol. 2006;26(4):1549–1557.

95. Hiemstra IH, Beijer MR, Veninga H, et al. The identification and developmental requirements of colonic CD169(+) macrophages. Immunology. 2014;142(2):269–278.

96. Fuchs T, Puellmann K, Hahn M, et al. A second combinatorial immune receptor in monocytes/macrophages is based on the TCRgammadelta. Immunobiology. 2013;218(7):960–968.

97. Beham AW, Puellmann K, Laird R, et al. A TNF-regulated recombinatorial macrophage immune receptor implicated in granuloma formation in tuberculosis. PLoS Pathog. 2011;7(11):e1002375.

98. Ajuebor MN, Das AM, Virag L, Flower RJ, Szabo C, Perretti M. Role of resident peritoneal macrophages and mast cells in chemokine production and neutrophil migration in acute inflammation: evidence for an inhibitory loop involving endogenous IL-10. J Immunol. 1999;162(3):1685–1691.

99. Soehnlein O, Lindbom L. Phagocyte partnership during the onset and resolution of inflammation. Nat Rev Immunol. 2010;10(6):427–439.

100. Rosales C. Neutrophil: A Cell with Many Roles in Inflammation or Several Cell Types? Front Physiol. 2018;9:113.

101. Evans JG, Chavez-Rueda KA, Eddaoudi A, et al. Novel suppressive function of transitional 2 B cells in experimental arthritis. J Immunol. 2007;178(12):7868–7878.

102. van de Veen W, Stanic B, Wirz OF, Jansen K, Globinska A, Akdis M. Role of regulatory B cells in immune tolerance to allergens and beyond. J Allergy Clin Immunol. 2016;138(3):654–665.

103. Blair PA, Norena LY, Flores-Borja F, et al. CD19(+)CD24(hi)CD38(hi) B cells exhibit regulatory capacity in healthy individuals but are functionally impaired in systemic Lupus Erythematosus patients. Immunity. 2010;32(1):129–140.

104. Smith TD, Tse MJ, Read EL, Liu WF. Regulation of macrophage polarization and plasticity by complex activation signals. Integr Biol (Camb). 2016;8(9):946–955.

105. Krishnamoorthy N, Oriss TB, Paglia M, et al. Activation of c-Kit in dendritic cells regulates T helper cell differentiation and allergic asthma. Nat Med. 2008;14(5):565–573.

106. Rodriguez-Cruz A, Vesin D, Ramon-Luing L, et al. CD3(+) Macrophages Deliver Proinflammatory Cytokines by a CD3- and Transmembrane TNF-Dependent Pathway and Are Increased at the BCG-Infection Site. Front Immunol. 2019;10:2550.

107. Strauss L, Mahmoud MAA, Weaver JD, et al. Targeted deletion of PD-1 in myeloid cells induces antitumor immunity. Sci Immunol. 2020;5(43).

108. Brandum EP, Jorgensen AS, Rosenkilde MM, Hjorto GM. Dendritic Cells and CCR7 Expression: An Important Factor for Autoimmune Diseases, Chronic Inflammation, and Cancer. Int J Mol Sci. 2021;22(15).

109. Forster R, Davalos-Misslitz AC, Rot A. CCR7 and its ligands: balancing immunity and tolerance. Nat Rev Immunol. 2008;8(5):362–371.

110. Strunk T, Temming P, Gembruch U, Reiss I, Bucsky P, Schultz C. Differential maturation of the innate immune response in human fetuses. Pediatr Res. 2004;56(2):219–226.

111. Filardy AA, Ferreira JRM, Rezende RM, Kelsall BL, Oliveira RP. The intestinal microenvironment shapes macrophage and dendritic cell identity and function. Immunol Lett. 2023;253:41–53.

112. Faria AM, Weiner HL. Oral tolerance. Immunol Rev. 2005;206:232–259.

113. Bhatta R, Han J, Liu Y, et al. Metabolic tagging of extracellular vesicles and development of enhanced extracellular vesicle based cancer vaccines. Nat Commun. 2023;14(1):8047.

114. Sabanovic B, Piva F, Cecati M, Giulietti M. Promising Extracellular Vesicle-Based Vaccines against Viruses, Including SARS-CoV-2. Biology (Basel). 2021;10(2).

